# Biophysical Models of PAR Cluster Transport by Cortical Flow in *C. elegans* Early Embryogenesis

**DOI:** 10.1101/2021.06.28.450200

**Authors:** Cole Zmurchok, William R. Holmes

## Abstract

The clustering of membrane-bound proteins facilitates their transport by cortical actin flow in early *Caenorhabditis elegans* embryo cell polarity. PAR-3 clustering is critical for this process, yet the biophysical processes that couple protein clusters to cortical flow remain unknown. We develop a discrete, stochastic agent-based model of protein clustering and test four hypothetical models for how clusters may interact with the flow. Results show that the canonical way to assess transport characteristics from single particle tracking data used thus far in this area, the Péclet number, is insufficient to distinguish these hypotheses and that all models can account for transport characteristics quantified by this measure. However, using this model, we demonstrate that these different cluster-cortex interactions may be distinguished using a different metric, namely, the scalar projection of cluster displacement on to the flow displacement vector. Our results thus provide a testable way to use existing single particle tracking data to test how endogenous protein clusters may interact with the cortical flow to localize during polarity establishment. To facilitate this investigation, we also develop both improved simulation and semi-analytic methodologies to quantify motion summary statistics (e.g., Péclet number and scalar projection) for these stochastic models as a function of biophysical parameters.

## 1 Introduction

Clustering of cell surface proteins is an important step in generating cell polarity. The PAR polarity proteins are a conserved network of proteins that localize asymmetrically in polarized cells that localize to the cell membrane during polarity establishment where they can be influenced by the flowing actin cortex (Goldstein and Macara, 2007). In the *C. elegans* single cell embryo, anterior PARs (aPARs) and posterior PARs localize to the anterior and posterior of the cell to organize the developing embryo. This organization results from a sperm-derived cue via biochemical interactions and due to the transport of proteins by the flowing actin cortex (Lang and Munro, 2017). The biophysical interactions between the cortex and membrane-bound proteins that facilitate this transport however remain unclear. Here we will use computational modeling to encode a range of different hypothesized types of protein-cortex interactions and demonstrate using these models that single particle tracking data quantified using an appropriate transport metric be used to distinguish these mechanisms from existing data modalities.

One of the aPAR proteins, PAR-3, is a scaffold protein that oligomerizes to form clusters (Harris, 2017; Thompson, 2021). In turn, PAR-3 clusters form complexes with other PAR proteins such as PAR-6, the kinase PKC-3 (aPKC) and small GTPase CDC-42 that are responsible for the core molecular interactions leading to polarization (Goehring, 2014; Lang and Munro, 2017; Sailer et al., 2015). Recent experimental work, summarized by Munro (2017), reveals a key role for clustering in the polarization process— PAR-3 is not transported by cortical flow and normal polarization does not occur without clustering (Dickinson et al., 2017; Rodriguez et al., 2017; Wang et al., 2017). These experimental investigations also revealed that the cortical residence times of clusters and their persistence of motion both increase with cluster size, raising the more specific question of how larger clusters are more effectively transported. Chang and Dickinson (2021) engineered PAR-3 clusters of various sizes and found that clusters of size three (i.e., consisting of three PAR-3 monomers) or larger are the minimum size required transport by flow to obtain polarity.

The primary data currently being used to assess protein transport in this area is single particle tracking coupled with fluorescent determination of protein cluster size. With this type of data in mind, protein cluster motion can be viewed as a size-dependent drift, diffusion process. The question then becomes, how do these clusters interact with the cortex to produce advection and how is that interaction modulated by cluster size. One broad hypothesis is that the clusters are subject to a size dependent drag force by the underlying flow. Another is that they (un)bind to the cortex with either a size-dependent or size-independent affinity. New experimental observations by Chang and Dickinson (2021) suggest that viscous forces on larger clusters are responsible for their transport and that PAR-3 clusters may not directly bind the actin cortex (since the clusters do not appear to co-localize with actin). Others note that these hypotheses remain broadly untested. For example, Gubieda et al. (2020) explain that

> it is not known exactly how clustering stabilizes PAR-3 at the membrane, but we can speculate that the coalescence of multiple membrane-binding domains could synergize to increase avidity for the membrane (Lemmon, 2008), and in a similar way, multiple membrane contact sites might increase resistance and prevent the lateral diffusion of the cluster. In addition, cluster size alone may also restrict diffusion or alter the ability of clusters to associate with or be corralled by features in the membrane or the cortex. More work is needed to dissect the precise mechanistic basis of flow-sensing by clusters,

and Illukkumbura et al. (2020) suggest that

> the precise mechanisms underlying rearward flow can differ and remain unclear in some systems because of the multi-faceted nature of clustering.

Here, we use mathematical modeling to explore the various facets of clustering to understand how are PAR-3 clusters are transported by the flowing cortex.

To investigate how clusters are transported by the flowing cortex, we developed, simulated, and analyzed a family of mathematical models for clustering and transport. These agent-based models (ABM) account for cluster growing and shrinking dynamics, binding and unbinding to the cortex, and a physics-based model for the cluster’s motion. We study four possible biophysical hypotheses for how the clusters may interact with the cortical flow, encoded as four different models. Where possible, we draw values for parameters in these models from literature. Where not, we either constrain them fitting the models to Péclet number observation data in Dickinson et al. (2017) (though we do not have access to the data itself, only figures from the article) or screen over indeterminable parameter ranges to ensure robustness of conclusions. We do note that the scope of our investigation is limited to understanding the interaction between these protein clusters and the cortical flow, and how that generates directed transport.

In this sense, our question and approach are complementary to many of the other modeling investigations of PAR polarity in the single-cell *C. elegans* embryo. A number of models have been developed to understand properties of the PAR signaling network. These models are typically partial differential equation models that capture the diffusion, biochemical reactions, and transport due to flow in the system (Aras et al., 2018; Dawes and Iron, 2013; Geßele et al., 2020; Goehring et al., 2011; Gross et al., 2018; Kravtsova and Dawes, 2014; Seirin-Lee et al., 2020; Wigbers et al., 2020). Goehring et al. (2011) for example found that a wave-pinning like mechanism (Holmes and Edelstein-Keshet, 2016; Holmes et al., 2012a,b, 2017; Jilkine and Edelstein-Keshet, 2011; Jilkine et al., 2007; Lin et al., 2012; Marée et al., 2006; Mata et al., 2013; Mori et al., 2008; Zmurchok and Holmes, 2020) may be responsible for stabilization of the anterior-posterior border. Other models focus on the events that follow polarization as the embryo further develops (Hubatsch et al., 2019). Dawes and Munro (2011) focused on the role that PAR-3 oligomerization has in the polarization process, finding that cluster may generate a bistable switch in the underlying kinetics (thus supporting polarized patterns) independent of protein cluster transport by flow. While these investigations focused primarily on understanding PAR signaling kinetics, our work is most similar to other investigations that seek to understand how proteins that switch state (i.e., with different diffusion coefficients in each state) can generate protein gradients (Bressloff et al., 2019; Wu et al., 2018) or investigations into the stochastic motion of PomX and PomY clusters in bacterial division (Bergeler and Frey, 2018; Kober et al., 2019).

The outcomes of our modeling study are threefold. First, we have developed a family of mathematical models that codify biophysically reasonable hypotheses for how the dynamics of these clusters and their potential interactions with the cortex influence their flow. We have also constrained parameters of these models to reasonable ranges. Second, we have identified a specific analysis metric that may be used to assess which of these hypotheses is most likely from existing single particle tracking data (Dickinson et al., 2017) or similar without the need to introduce engineered forms of these proteins. Third, we have provided new computational and semi-analytic approaches to calculate motion statistics (Péclet number and scalar projection) that simplify and speed the analysis of these models. While we cannot conclude which of the mechanisms described in these models is responsible for cluster transport in the developing *C. elegans* embryo, this combination of models and analysis approach coupled with appropriate data may provide a means to do so.

## 2 Methods

We model PAR-3 protein clusters using a discrete, stochastic agent-based model (ABM). Our purpose is to use this modeling framework to encode and test several hypotheses regarding the biophysical interactions between protein cluster and the flowing actin cortex. Ultimately, we will compare model predictions to experimental data from Dickinson et al. (2017). In this section, we outline our computational approach, describe the different biological hypotheses (Models 1 through 4) encoded with this approach, and describe how we estimated the model parameters.

We assume that PAR-3 protein clusters (henceforth clusters), illustrated in Figure 1, grow via the addition of monomers, and that, being bound to the cell membrane, can move in two-dimensions (panel B). From a force-balance equation, we obtain a stochastic-differential equation (SDE) for the motion of each cluster, with the drift and diffusion terms that depend explicitly on the cluster size and the specific biological hypothesis under consideration. Using biophysically realistic parameter ranges, we simulate cluster dynamics in the agent-based simulation and develop a faster and more efficient simulation method based on Monte-Carlo (MC) sampling from appropriate distributions. Finally, we adopt model outputs such as the Péclet Number (as defined by Dickinson et al. (2017)) and the scalar projection of the cluster’s displacement in order to compare the transport of our simulated clusters with experimental data.

**Figure 1:**
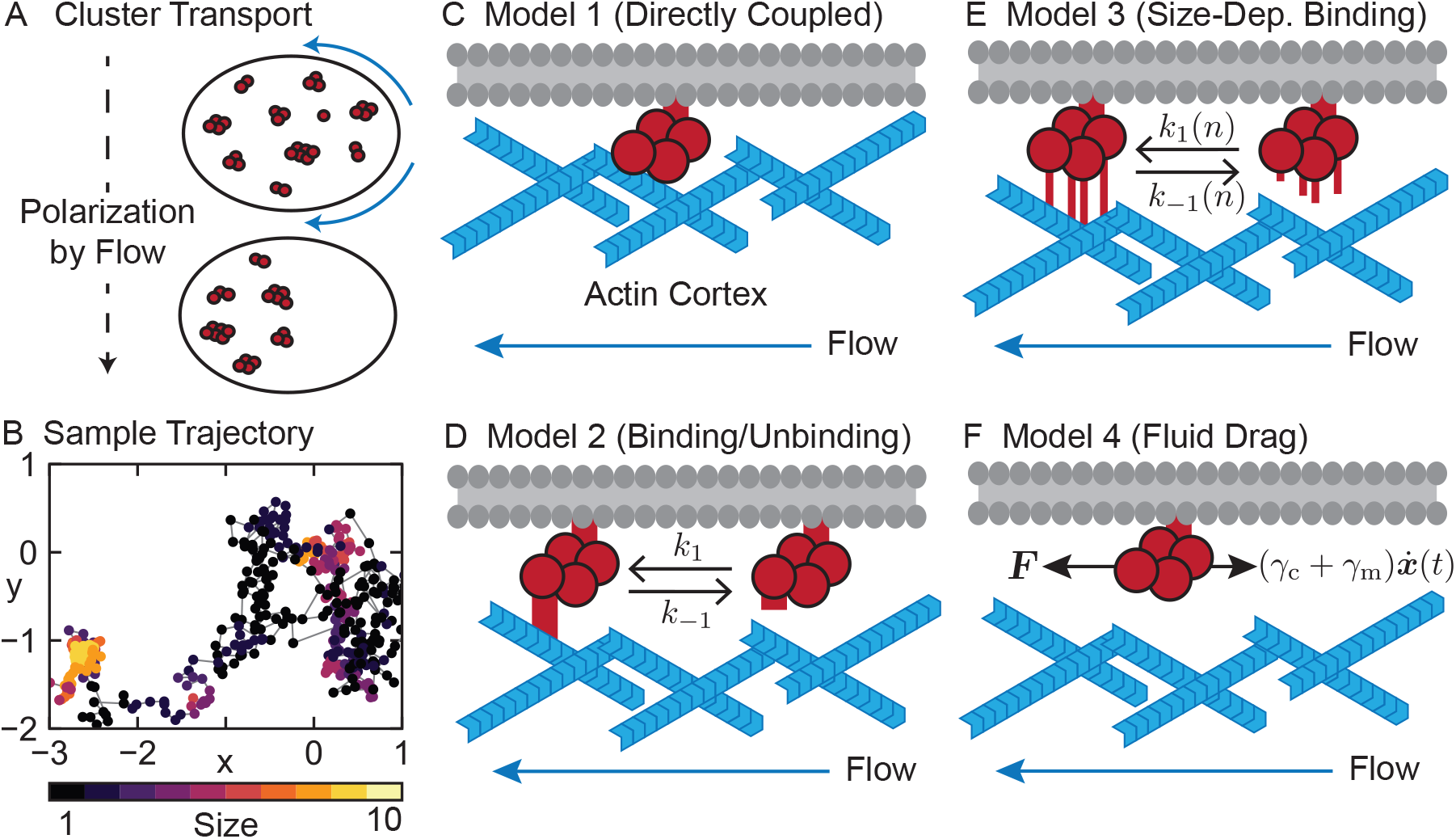
Cluster Transport by Flow, Sample Trajectory, and Model Cartoons. A. PAR-3 clusters are transported by cortical flow (blue arrows) in the *C. elegans* embryo (adapted from Munro (2017)). Each cluster is assumed to be bound to the cell membrane and gains or loses monomers with constant rates *k*_+_ and *k*_−_, respectively, and diffuses in the membrane. B. Sample trajectory of a diffusing cluster with ***x***(0) = (0, 0) and no coupling to cortical flow. Cluster position and size is shown every Δ*t* = 1 s. Since the diffusion coefficient decreases with cluster size, the cluster appears to move less when it is larger. C-F. Hypothesized models for cluster-cortex interactions. In Model 1 (panel C), we assume that clusters are always bound to the flowing actin cortex and move with the flow velocity. In Model 2 (panel D), we assume that clusters stochastically bind and unbind to the cortex with size-independent rates *k*_1_ and *k*_−1_, while in Model 3 (panel E), we assume that clusters bind and unbind with size-dependent rates *k*_1_(*n*) and *k*_−1_(*n*). In Model 4 (panel F), we assume that a drag force ***F*** generated by the flowing actin cortex opposes the drag that the cluster experiences from the environment and cell membrane.

### 2.1 Cluster Dynamics Model

We model each cluster as an independent agent described by three random variables: ***x***(*t*) ∈ ℝ^2^ gives the cluster’s position, *n*(*t*) = 1, 2, 3, … describes the integer number of monomers in the cluster, and *b*(*t*) ∈ {−1, 1} describes whether the cluster is unbound (−1) or bound (1) to the cortex at time *t*. To be clear, we differentiate between the membrane and the flowing cortex here. All clusters are assumed to be bound to the cell membrane at all times since those that are not would not be observed using the experimental quantification motivating this article (Dickinson et al., 2017). From here on, (un)bound refers to the interaction between the clusters and the cortex. The form of this interaction is precisely what we are investigating here.

#### Size Dynamics

To describe the change in cluster sizes we adopt a simplified version of the simple polymerization model (Edelstein-Keshet and Ermentrout, 1998) used to model the polymerization of actin filaments (also described in Chapter 4.1 of Bressloff (2014)). Monomers are added to the cluster with rate *k*_+_ and are removed with rate *k*_−_. The probability *p*_*n*_(*t*) that the cluster is of size *n* at time *t* is described by:

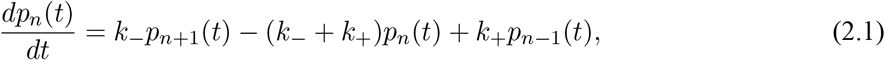

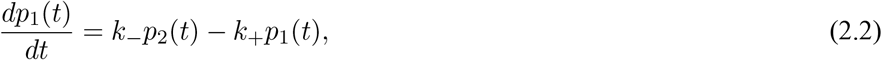

with the normalization condition 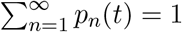. A stationary solution satisfies

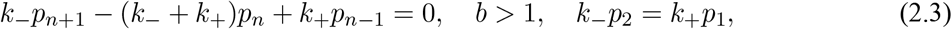

motivating the ansatz *p*_*n*_ = *Cλ*^*n*^. This results in *λ* = 0, 1, or *λ* = *k*_+_/*k*_−_. To satisfy the normalization condition with nontrivial cluster dynamics, we must have *λ* = *k*_+_/*k*_−_ < 1. In this case *C* = (1 − *k*_+_/*k*_−_)/(*k*_+_/*k*_−_). Thus, the steady-state clustering dynamics satisfy

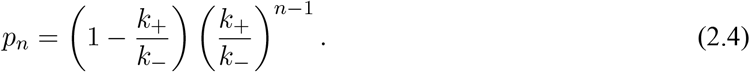

Note that we have assumed that the concentration of monomers is unlimited in this scenario and have neglected higher order growth and decay events (e.g., the addition and subtraction of oligomers of any size).

#### Binding Dynamics

We assume that the clusters can bind (*b*(*t*) = 1) and unbind (*b*(*t*) = −1) to the cortex with reaction rates *k*_1_ and *k*_−1_, respectively. Depending on the specific biophysical hypothesis for how the cluster interacts with the flowing actin cortex, we may consider the limit where *k*_1_ → ∞ and *k*_−1_ → 0, or allow the binding and unbinding rates to depend on the cluster size *n*(*t*). These assumptions will be clarified in the next section.

With *p*_*b*_(*t*) defined as the probability that the cluster is bound to the actin cortex at time *t*, we obtain the following master equation:

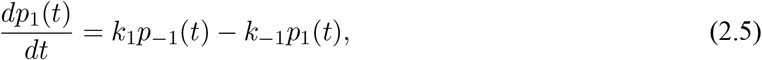

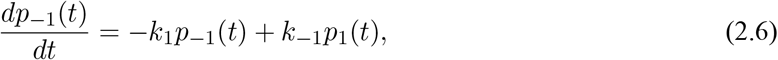

with stationary solution

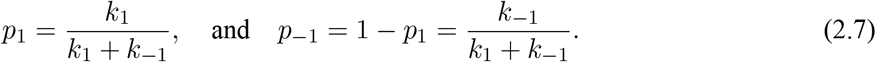

#### Position Dynamics

Newton’s equations for the cluster, neglecting inertial forces, give

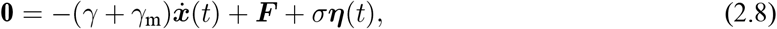

where ***x***(*t*) = (*x*_1_(*t*), *x*_2_(*t*)) ∈ ℝ^2^ is the stochastic position of the cluster at time *t*, ***F*** = (*F*_1_, *F*_2_) is an external force (e.g., from the flowing actin cortex), *γ* and *γ*_m_ are drag coefficients, *σ* is the noise level, and ***η***(*t*) = (*η*_1_(*t*), *η*_2_(*t*)) is Gaussian white noise. There are two potential sources of drag on the cluster: drag on the cluster from the membrane-binding site with drag coefficient *γ*_*m*_, and drag from the cellular environment with drag coefficient *γ*. For simplicity, we assume Stokes’ Law:

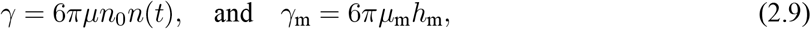

where the cluster’s radius is assumed to be linearly proportional to the number of monomers *n*(*t*) in the cluster (*n*_0_ being the size of each monomer) and *μ* is the viscosity. We also assume that the membrane-binding site drag can also be described using Stokes’ Law, with radius given by the membrane height *h*_m_, and membrane viscosity *μ*_m_. This drag does not depend on cluster size since it is assumed the cluster is anchored to the membrane by a separate domain. Note that we adopt this simple model of drag given the lack of specific structural data on PAR-3 complexes (to our knowledge). More discussion of these drag coefficients can be found in the methods section discussing parameters. From equation (2.8), we obtain a stochastic differential equation (SDE) for the position of the cluster:

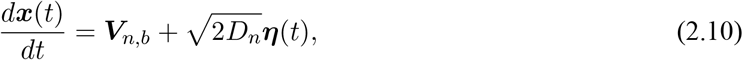

where the drift coefficient ***V***_*n,b*_ depends on the cluster’s size *n*(*t*) and bound state *b*(*t*) through the external force ***F*** and size-dependent viscous drag:

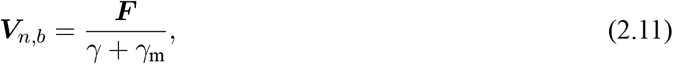

and the diffusion coefficient *D*_*n*_ depends on the cluster’s size *n*(*t*) through the size-dependent viscous drag:

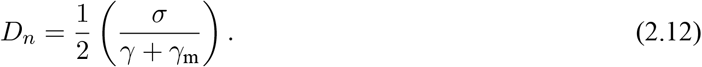

Note that the diffusion and drag coefficients are jointly defined to satisfy the relevant fluctuation-dissipation relation.

The full cluster dynamics can then be described through a chemical master equation for the probability, *P* (***x***, *n, b, t*), that a cluster is at location ***x***, is of size *n*, and has binding state *b* at time *t*:

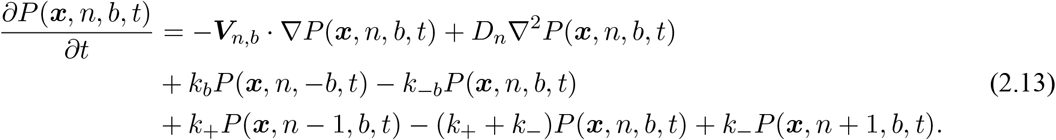

We provide this description for those who are familiar with such processes. However, in practice we either directly simulate cluster dynamics or approximate their statistical properties in some other way and do not directly work with the master equation.

### 2.2 Models 1–4: Interaction with the Cortical Flow

We study four possible biophysical hypotheses for how the cluster may interact with the cortical flow to give rise to cluster transport. Our list is not exhaustive but seeks to capture the main ideas of how such clustering may enhance transport. We assume that the clusters remain bound to the cell membrane throughout the transport process. To organize these hypotheses, we developed a series of four models. These models are illustrated in Figure 1 C–F. Note that the flowing actin cortex is described by a velocity field ***υ***_c_.

1. **Model 1: Directly Coupled**. We assume that the cluster is directly coupled to the flowing cortex, so that the cluster’s velocity (in the absence of noise) exactly matches the cortical velocity, ***υ***_c_. In this case, the external force imposed by the cortex on the cluster is ***F*** = (*γ* + *γ*_*m*_)***υ***_c_ so the drift term is ***V***_*n,b*_ = ***υ***_c_ for all *n* and *b*.
2. **Model 2: Binding/Unbinding**. We assume that the cluster can bind and unbind to the flowing cortex with reaction rates *k*_1_ and *k*_−1_ respectively, and that the cluster moves with the cortex velocity when bound. In this case, the external force imposed by the cluster is 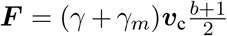. In other words, the drift coefficient switches between **0**and ***υ***_c_ as the cluster binds and unbinds from the cortex.
3. **Model 3: Size-Dependent Binding/Unbinding**. We make the same assumptions as in Model 2; however, we assume that the cluster size can influence the binding kinetics. This assumption seeks to model the hypothesis that larger clusters may have multiple binding domains that could synergize coupling to the flowing cortex (e.g., increasing the avidity for the cortex). Thus, we assume that the binding rate increases linearly with cluster size:

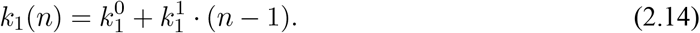 This effectively ensures that larger clusters remain bound to the flowing cortex for a longer amount of time. Multiple functional forms could be envisioned for this relation or alternatively the size dependence could effect the off rate. We use this size dependence for simplicity to compare against these other different hypotheses.
4. **Model 4: Fluid Drag**. Our fourth hypothesis is substantively different from the binding/unbinding models in that we assume that the flowing cortex exerts a drag force on the cluster that is opposed by the drag from the membrane and that there is no overall drag in the system. Thus, instead of equation (2.8), we have

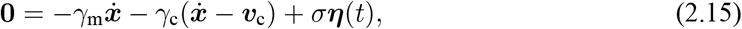

where *γ*_*c*_ = 6*πμn*_0_*n*(*t*) is defined analogously to *γ*. Under these assumptions, the drift term directly depends on the cluster size *n*:

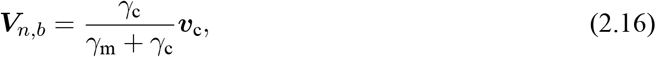

and the diffusion coefficient is:

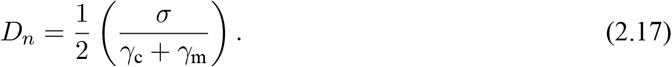

There are similarities and differences between these models. Model 1 can be thought of as taking a limit of model 2 when *k*_1_ → ∞ and *k*_−1_ → 0, so that the cluster velocity always matches the cortical velocity whereas in model 2, we expect the cluster’s velocity to match the cortex velocity for a proportion, 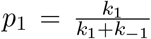, of the time it is moving. In models 1–3, the drift coefficient is size-independent, and size dependence only enters Model 3 through (un)binding rates. This is different from model 4, where both the drift and diffusion coefficients depend on cluster size— equation (2.16) specifies that larger clusters are more affected by the flowing cortex since *V*_*n,b*_ is a saturating function in *n*. Note that we consider two possible sources of size-dependent effects in our models: (1) through the diffusion coefficient *D*_*n*_ and (2) through the binding/unbinding dynamics (or fluid drag force) that give rise to the drift coefficient *V*_*n,b*_. This will be important because, as we will see, the size effects on net drift rate can markedly distinguish these hypotheses while the size-dependent diffusivities are essentially the same across models.

### 2.3 Quantifying Cluster Transport

The primary data in this area quantifies cluster sizes (via fluorescence brightness) and tracks cluster motions as a function of rough size (Dickinson et al., 2017). While we do not have access to this data, we will analyze these models with quantitative measures that could be applied to this type of data.

Dickinson et al. (2017) quantified the size-dependent cluster transport by calculating an apparent Péclet number. To compare our simulation results with experimental data, we adopt the same definition:

> For each particle at each time step, we calculated a vector for the motion due to advection (which is equal to the local cortical flow...) and a vector for the motion due to diffusion (which we estimate as the total particle displacement minus the advective motion). The apparent Péclet number for that time step is simply the ratio of the lengths of the advection vector and the diffusion vector.

We illustrate the definition of the Péclet number in Figure 2A. Define the cluster’s displacement over the time step of length Δ*t* as ***r*** = ***x***(*t*+Δ*t*) − ***x***(*t*). The motion due to cortical flow in this time step is ***υ*** = ***υ***_c_Δ*t* and so the motion due to diffusion is ***d*** = ***r − υ***. The Péclet number for that time step is defined as the ratio of the lengths of these vectors:

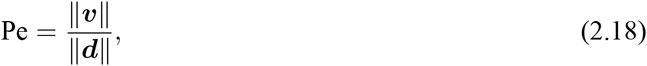

provided that ***d*** ≠ **0**. Results below will show that Péclet number is ineffective at distinguishing these models. Thus we introduce another metric of cluster transport that compare the cluster’s motion relative to the flow (as opposed to the cluster’s motion relative to the noise). For this second metric, we calculate the scalar projection of the cluster’s displacement ***r*** onto the cortical flow ***υ***:

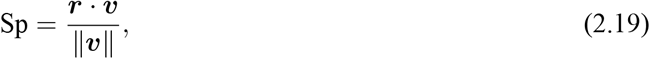

provided that υ ≠ **0**.

**Figure 2:**
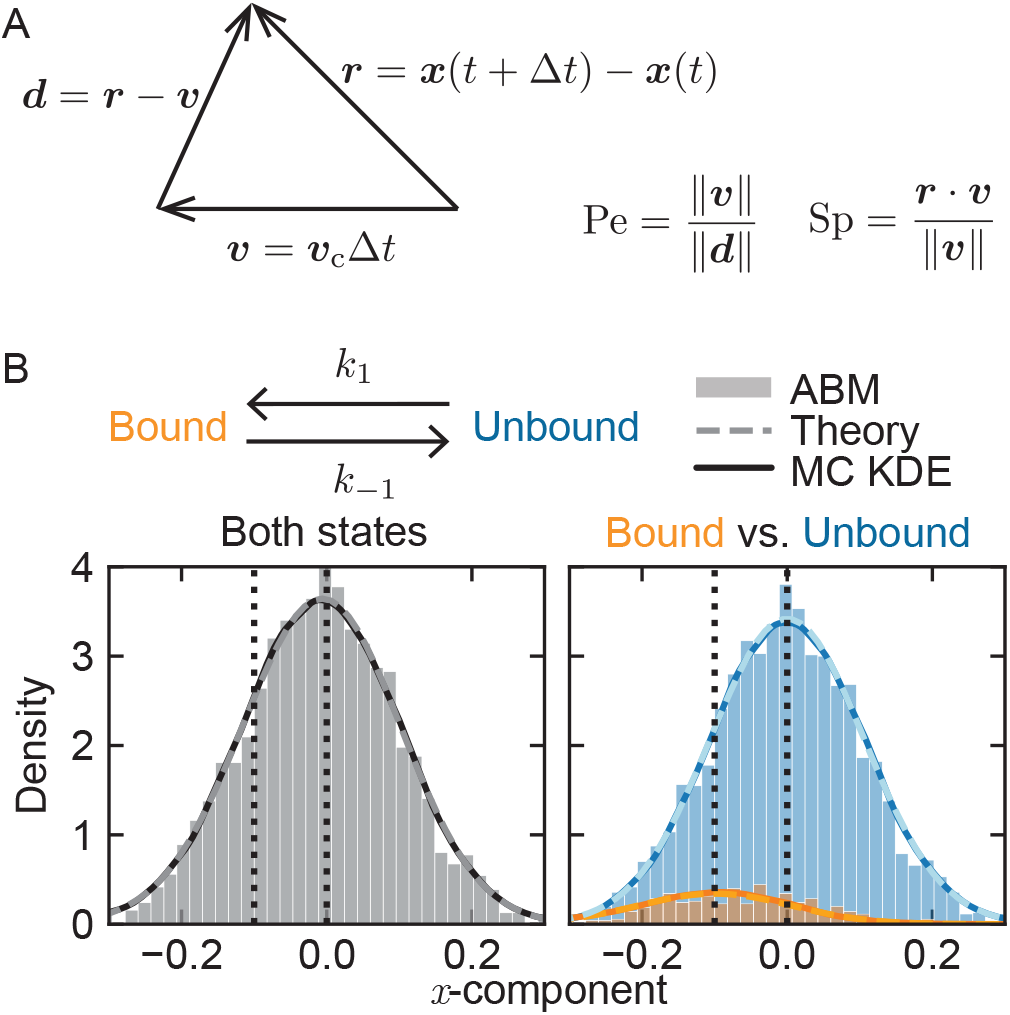
A. Metrics used to quantify cluster transport. Each cluster has displacement (in time Δ*t*) defined as ***r*** = ***x***(*t* +Δ*t*) − ***x***(*t*) that can be decomposed into an advective component from cortical flow ***υ*** = ***υ***_c_Δ*t* and a diffusive component ***d*** = ***r − υ***. The Péclet number, Pe, is defined as the ratio of the lengths of the advective component and the diffusive component. The scalar projection Sp, is given by the scalar projection of the displacement onto the advective component. B. Probability densities of the *x*-component of cluster displacements, ***r***_1_ (see panel A), obtained from the simulations of the agent-based model (ABM; histograms) match the kernel density estimates of distributions obtained from Monte-Carlo (MC) sampling (solid lines) from quasi-steady distributions (dashed lines; equations (2.25)–(2.27)). The distribution for clusters of size *n* = 4 in both bound and unbound states is shown in the left panel whereas the distributions in the bound and unbound states are shown separately in the right panel. Vertical lines at 0 and −0.1 illustrate how unbound clusters have mean displacement of 0 while bound clusters have mean displacement equal to the cortical flow speed (0.1 *μ*m/s; we assume that flow is always in the negative *x*-direction). 100 cells were simulated until time *T* = 500 s, i.e., 50,000 total displacements) from clusters of size *n* = 4 using MC sampling while 10000 cells were simulated until time *T* = 500 s using the ABM with *σ* = 0.01, *μ* = 0.1, *k*_1_ = 0.1, *k*_−1_ = 1. Supplemental Figures 2–5 illustrate how the displacement distributions obtained from the ABM and MC sampling methods agree over all cluster sizes.

### 2.4 Stochastic Simulation Methods

To numerically simulate the cluster dynamics, we use two complementary approaches. The first is to use a “naive” stochastic simulation algorithm as in Chapter 1 of Erban and Chapman (2019) to generate a stochastic trajectory of the cluster dynamics (position, growth/decay, and binding/unbinding). We refer to this method as the “agent-based model” (ABM). The second method is more efficient and is based on Monte-Carlo (MC) sampling. In the MC sampling method we do not track the cluster’s position over time, but instead generate many cluster displacements ***r*** by sampling from the appropriate statistical distributions. In this section, we describe both approaches.

#### 2.4.1 Agent-Based Model

We simulate *N* non-interacting clusters from time *t* = 0 s to *t* = *T* = 500 s approximating the 8 minutes during which the *C. elegans* embyro is in the establishment phase and the PAR proteins are clustered (Dickinson et al., 2017). We initialize each cluster at ***x***(0) = (0, 0) with *n*(0) ~ *U* (1, 10) (randomly sampled from a uniform distribution of integers between 1 and 10), and *b*(0) = −1. At each time step Δ*t*_ABM_ = 0.001 s, we determine **V**_*n,b*_ and *D*_*n*_ from equations (2.11) and (2.12) (or (2.16) and (2.17) in Model 4). Next, we update the position of the cluster:

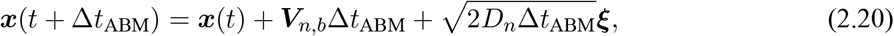

where ***ξ*** = (*ξ*_1_, *ξ*_2_) has components *ξ*_*i*_ that are independent samples obtained from a standard normal distribution. We assume that Δ*t*_ABM_ is sufficiently small such that at most one reaction occurs (adding or subtracting a monomer, binding or unbinding to the cortex) to occur. To determine which reaction occurs, we first label the reactions: *R*_*k*_, *k* = 1, … , 4 where *k* = 1 corresponds to adding a monomer, *k* = 2 to subtracting a monomer, *k* = 3 to binding to the cortex, and *k* = 4 to unbinding from the cortex. Each reaction has probability *p*_*k*_ given by the product of the reaction rate and the time step Δ*t*_ABM_: *p*_1_ = *k*_+_Δ*t*_ABM_, *p*_2_ = *k*_−_Δ*t*_ABM_, *p*_3_ = *k*_1_Δ*t*_ABM_, and *p*_4_ = *k*_−1_Δ*t*_ABM_. To determine which reaction occurs, we draw a random number *r* ~ *U* (0, 1) and find *k* such that

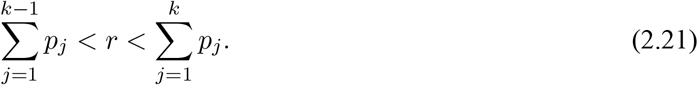

If no *k* satisfies this condition, then no reaction occurs in the time step. We then determine *n*(*t* + Δ*t*_ABM_) and *b*(*t* + Δ*t*_ABM_) as depending on which reaction occurs:

1. If *R*_1_ occurs, then *n*(*t* + Δ*t*_ABM_) = *n*(*t*) + 1 and *b*(*t* + Δ*t*_ABM_) = *b*(*t*).
2. If *R*_2_ occurs, then *n*(*t* + Δ*t*_ABM_) = *n*(*t*) − 1 (only if *n*(*t*) > 1) and *b*(*t* + Δ*t*_ABM_) = *b*(*t*).
3. If *R*_3_ occurs, then *n*(*t* + Δ*t*_ABM_) = *n*(*t*) and *b*(*t* + Δ*t*_ABM_) = 1.
4. If *R*_4_ occurs, then *n*(*t* + Δ*t*_ABM_) = *n*(*t*) and *b*(*t* + Δ*t*_ABM_) = −1.
5. If no reaction occurs, then *n*(*t* + Δ*t*_ABM_) = *n*(*t*) and *b*(*t* + Δ*t*_ABM_) = *b*(*t*).

We save the data for analysis every Δ*t* = 1 second. The model is simulated with a smaller time step, however 1 second matches the imaging frequency of the motivating data and so we use it for analysis.

To ensure that the numerical implementation of the ABM is correct, we simulated *N* = 1000 clusters with *D*_*n*_ = 0.001 for all *n*, and with or without drift (***V***_*n,b*_ = ***υ***_c_ or **0**). We then calculated the mean-square displacement (MSD), the time spent as a cluster of each size, and the cluster size distribution at the end of the simulation. Our stochastic simulations match (Supplemental Figure 1) standard theory for MSD for particles moving due to diffusion with or without drift, the expected time spent as a cluster of size *n*, (1/*k*_+_ for clusters of size 1 and 1/(*k*_+_ + *k*_−_) for clusters of all other sizes), and the steady-state cluster size distribution predicted by equation (2.4).

#### 2.4.2 Monte-Carlo Sampling

Instead of simulating each cluster until time *T* using the computationally expensive ABM, we developed a more efficient Monte-Carlo (MC) sampling method. In practice, we only quantify cluster transport through the Péclet number and the scalar projection. Since these quantities do not depend on the cluster position but only the cluster displacements ***r*** (Figure 2A) it is sufficient to determine the displacement distributions to determine the transport characteristics.

As motivation for this approach, consider clusters of size *n*. The motion of these clusters is characterized by a diffusion coefficient *D*_*n*_ and a drift coefficient **V**_*n,b*_ that also depends on the binding state *b*. Rewriting the SDE (2.10) in differential form

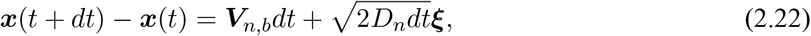

where ***ξ*** has independent standard normally distributed components, reveals that the displacements of clusters of size *n* in binding state *b* over time Δ*t* (denoted by ***r***_*n,b*_) are normally distributed with mean ***V***_*n,b*_Δ*t* and covariance 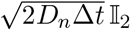 (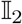 is the 2-by-2 identity matrix):

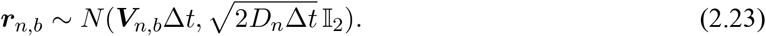

Overall, we exploit the fact that the SDE (2.10) relates the change in position of the cluster (displacement) to the drift and diffusion coefficients. Note that for models 1 and 4 the drift coefficient, ***V***_*n,b*_, does not depend on the binding state *b*. Thus, in order to simulate a cluster of size *n* for *T* /Δ*t* time steps using the MC sampling method in models 1 and 4, we draw *T* /Δ*t* samples from the appropriate distribution:

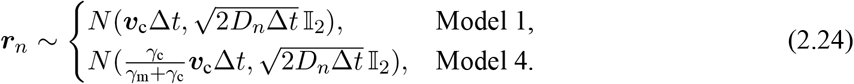

For models 2 and 3, however, we need to further simplify by averaging over the bound and unbound states. Assuming that the binding and unbinding process is in quasi-equilibrium, then using the stationary solution given by (2.7), we find that clusters of size *n* spend *p*_1_ proportion of time bound to the cortex and *p*_−1_ proportion of time unbound. Note that these probabilities depend on *n* in model 3 since *k*_1_ is a function of *n*, but are constant with respect to *n* in model 2. Thus, to simulate a cluster of size *n* for *T* /Δ*t* time steps, we first determine for how many of these time-steps the cluster is expected to be bound and how many unbound. We assume that each time step is independent (and that *T* /Δ*t* is an integer), and therefore find the number of time steps in the bound state to be binomially distributed with *T* /Δ*t* trials and probability *p*_1_:

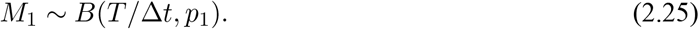

The number of time-steps spent in the unbound state, is easily calculated as *M*_−1_ = *T* /Δ*t − M*_1_. Thus, to simulate a cluster for *M* time steps in model 2 or 3, we draw *M*_1_ samples for the displacements while bound:

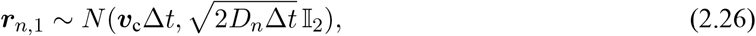

and then draw *M*_−1_ samples for the displacements while unbound:

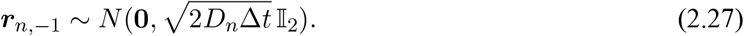

Note that in the MC sampling method we ignore the procession of time—we collect these displacements in a set. A true trajectory would have a ordered list of displacements corresponding to the changes in the cluster’s position, size, and binding state over time.

We validated the accuracy of the Monte-Carlo sampling approach by comparing the displacement distributions obtained through ABM simulations with those obtained through MC sampling, and comparing both of these distributions to their analytical forms (the probability density function for a multivariate normal distribution with given mean and covariance). A sample of this comparison is shown in Figure 2B for clusters of size *n* = 4. We show the probability densities of the *x*-component of the cluster displacements obtained from ABM simulations as a normalized histogram, kernel density estimates obtained from MC sampling (solid lines), and exact forms of the quasi-steady distributions (dashed lines). In the left panel, the bound and unbound displacements are aggregated while in the right panel the displacement distributions for the bound and unbound state are shown separately. Supplemental Figures 2–5 show the agreement between the displacement distributions obtained between ABM simulations and MC sampling method across all cluster sizes.

One advantage of the MC sampling method over the ABM is that large clusters are relatively rare. Given the biologically relevant parameter values for *k*_+_ and *k*_−_, the probability that a cluster is a large size (e.g., 8, 9, or 10) is very small (see Supplemental Figure 1C). Thus, in order to obtain reasonable statistics for the transport of clusters when they are large, it is necessary to run the ABM for a large number of simulations for sufficiently long time to collect enough transport data for these large clusters. In the MC sampling method, however, it is straightforward to sample a sufficient number of displacements for each size and determine the expected displacement distributions. Since we are only interested in broad transport metrics such as the Péclet number and scalar projection, we use the MC sampling method for the remainder of the paper.

### 2.5 Parameter Estimation

We estimated model parameters from literature wherever possible, and performed a sweep over ranges of key parameters in order to assess the transport of clusters. In this section, we describe the model parameters, parameter sweep, and summarize the model and output variables, parameters used in simulations, and numerical parameters in Table 1.

**Table 1:**
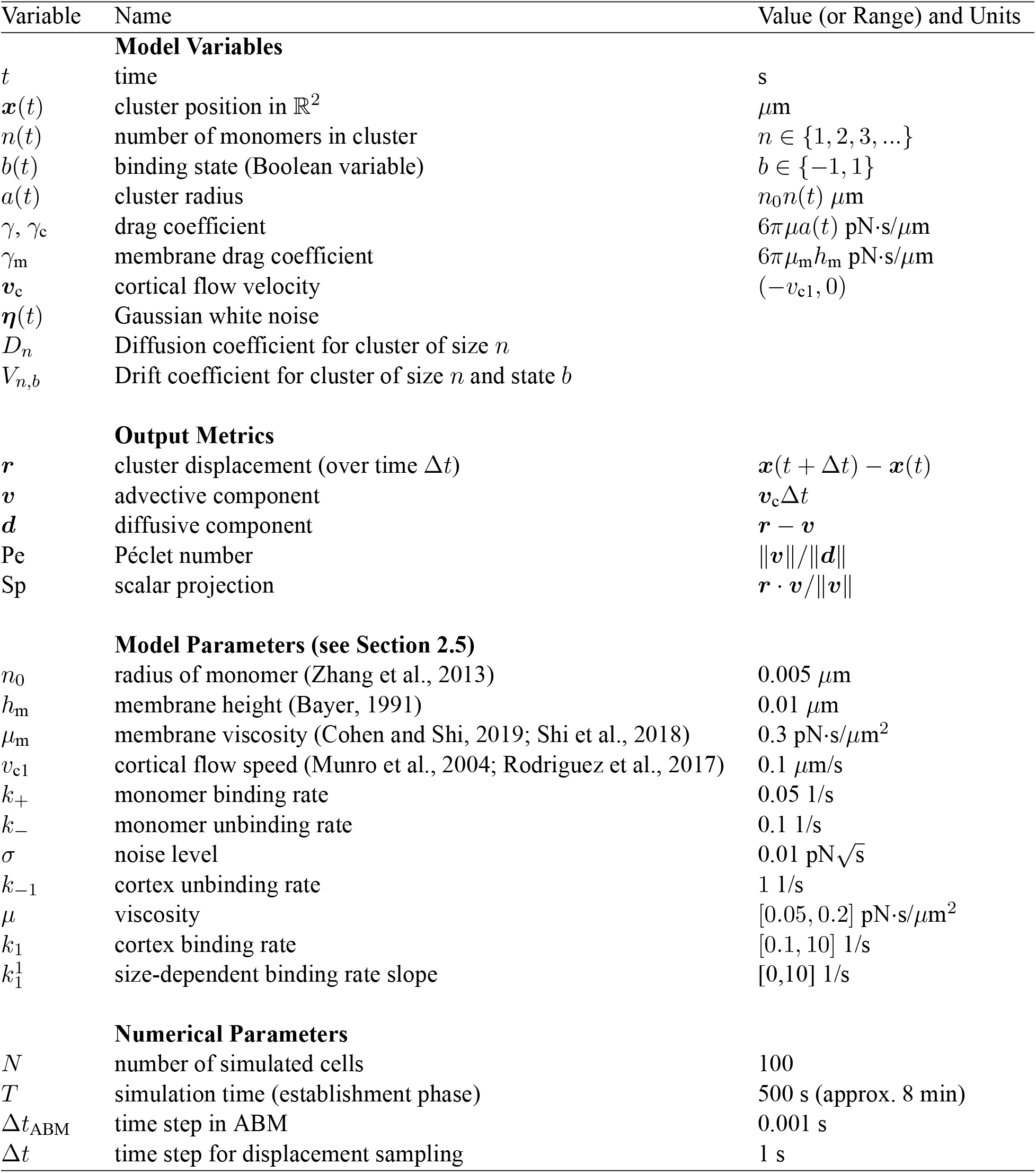
Summary of model variables, output metrics and parameters.

#### 2.5.1 Parameters from Literature

We use values obtained from literature for several of the model parameters. We use micrometers (*μ*m) for space, seconds (s) for time, and pico-Newtons (pN) for force. PAR-3 monomers are approximately 5 nm across (see Figure 2 of Zhang et al. (2013) and Figure 1 of Harris (2017)), so we set *n*_0_ = 0.005 *μ*m. We assume that the cell membrane is homogeneous and is 10 nm thick (*h*_m_ = 0.01 *μ*m, (Bayer, 1991)). We use the two-dimensional membrane viscosity estimates (0.003 pN·s/*μ*m) provided by Cohen and Shi (2019); Shi et al. (2018), but convert this viscosity to the appropriate dimensions (a 3D viscosity), by dividing by the cell membrane height: *μ*_m_ = 0.3 pN·s/*μ*m^2^. The cortical flow speed has been measured in *C. elegans* embryos by Munro et al. (2004) and by Rodriguez et al. (2017). Using the flow speed value of 7.66 ± 1.0 *μ*m/min (Rodriguez et al., 2017), we set *υ*_c1_ = 0.1 *μ*m/s (7.66/60 ≈ 0.13). We also assume that the flow direction is always in the negative *x*-direction, and thus set ***υ***_c_ = (−*υ*_c1_, 0). We justify this assumption by noting that we are only interested in how the clusters are affected by flow instead of the overall transport of clusters in the embryo. Finally, we set *k*_+_ = 0.05 1/s and *k*_−_ = 1.0 1/s so that the distribution of cluster sizes qualitatively matches the distribution from Figure 3B in Dickinson et al. (2017) during the establishment phase. Recall that steady-state cluster sizes are exponentially distributed, see equation (2.4) (Supplemental Figure 1C). We also find that the expected time in each size state is on the order of seconds or tens of seconds (Supplemental Figure 1B) with these parameter values. Since we are only interested in how cluster size affects the transport of the clusters in the sense of local displacements instead of overall cluster distribution in the embryo (and we do not explicitly simulate protein cluster size in the MC sampling method), the precise values of *k*_+_ and *k*_−_ are less important with one caveat. If monomer addition and subtraction were faster than the 1 second analysis time scale we use, some of the output quantification and analyses here may be affected. However, this would affect the interpretation of data as well. Thus since we do not have direct data on the timescale of monomer addition and subtraction, we assume the rates are slower than the 1 second observation timescale.

**Figure 3:**
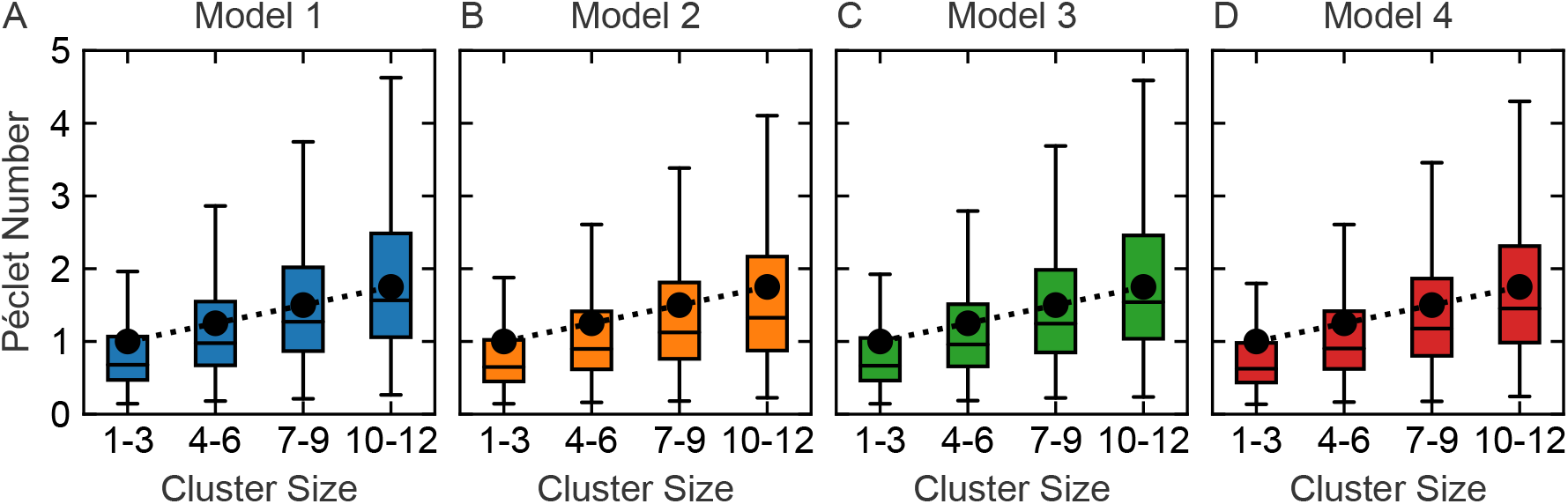
All models agree with observed trend in Péclet number in experimental data (panels A–D for models 1–4, respectively). Black points show mean Péclet numbers estimated from experimental observations of cluster transport from Figure 6H of Dickinson et al. (2017), while boxplots illustrate simulated Péclet number distributions obtained from MC sampling when cluster displacements are grouped into four size categories. Size-dependent diffusion, equation 2.12, is required to obtain an increase in mean Péclet number with cluster size. See Supplemental Figure 11 for Péclet number distributions when diffusion is size-independent. We simulated *N* = 100 cells with unique parameters and sampled 500 displacements for each cluster size for each model. Parameters: *σ* = 0.01, *μ* ~ *U*(0.05, 0.2), *k*_1_ ~ *U* (0.1, 10) (*k*_1_ = 5 is fixed in Model 3), *k*_−1_ = 1, and 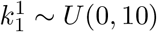.

#### 2.5.2 Parameter Sweep

We were unable to estimate the remaining parameters from literature. In order to constrain the noise level *σ*, the viscosity *μ*, the cortex unbinding rate *k*_−1_, the cortex binding rate *k*_1_, and the cortex size-dependent binding rate slope, we performed a parameter sweep. Based on Péclet number quantification from Dickinson et al. (2017), we assume the true mean Péclet number as a function of cluster size varies linearly between 1 and 2 for clusters of sizes 1 through 10 as a target for our parameter sweep. We sweep over parameter sets and only retain those that approximately satisfy this condition.

##### Noise Level and Viscosity

To start, we used model 1 as a test and sampled 500 displacements of each cluster size from 100 cells with *σ* ranging from 0.01 to 0.04 and *μ* ranging from 0.1 to 0.4. Results (Supplemental Figure 7) show that increasing the noise level increases the effect of diffusion (reducing the Péclet number) and increasing the viscosity reduces the effect of diffusion (increasing the Péclet number). We set *σ* = 0.01 since this gave the best results compared to our criteria when *μ* = 0.1. However, since we wish to account for any heterogeneity that may be present, we allowed *μ* to vary in a range. We chose *μ* ∈ [0.05, 0.2].

As a final check for biological realism, we estimated the diffusion length scale for freely diffusing clusters with *σ* = 0.01 and *μ* ∈ [0.05, 0.2]. The diffusion length scale for clusters of size *n* is given by 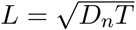 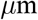 over time *T* = 500 s, and ranges from 0.6 to 2.6 *μ*m (Supplemental Figure 6) with these parameter values. These length scales provide an estimate that the clusters could travel only a few micrometers by diffusion alone, reasonable for membrane-diffusing particles of this size. Moreover, in the fluid drag model (model 4), we found that increasing the viscosity beyond *μ* = 0.20 gives unreasonably large Péclet numbers (Supplemental Figure 10).

##### Binding and Unbinding Rates

Here we again note that the ratio of these two rates is the main determinant of dynamics and we do not have sufficient data to parameterize the timescale of cortical binding / unbinding. Thus we first set *k*_−1_ = 1 1/s (effectively re-scaling time). Based on parameter screens, we find that *k*_1_ ∈ [0.1, 10] 1/s produce mean Péclet number that vary between 1 and 2 as a function of size (Supplemental Figure 8). In model 3, *k*_1_ increases with the cluster size via equation (2.14). We chose 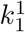 to vary between 0 and 10 in order to generate a range of size-dependent effects, but fixed 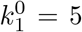 to reduce the number of parameters being varied in any given simulation. With 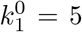 (fourth row Supplemental Figure 9), we found that the mean Péclet number across cluster sizes matched our criteria to increase from 1 and 2 as the cluster sizes increase.

#### 2.5.3 Parameters for Simulations

Rather than choose a single parameter set to analyze, we instead choose 100 parameter sets from the ranges listed above. In one sense, this allows us to generalize our analysis to the relevant region of parameter space. In another, each of these “parameter sets” can be thought of as an *in silico* cell and by doing this, we account for likely heterogeneity that would be naturally present. In summary, we fixed *σ* = 0.01, *k*_−1_ = 1, and allowed *μ*, *k*_1_, and 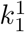 to vary over a range for our simulations. Using these parameter ranges, we generated *N* = 100 parameter sets uniformly sampling from the ranges: *μ* ~ *U* (0.05, 0.2), *k*_1_ ~ *U* (0.1, 10) (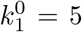 is fixed in Model 3), and 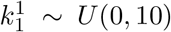. We think of these *N* = 100 parameter sets as representing 100 different embryos to be measured, and thus report our model outputs (mean Péclet number and scalar projection) as mean values plus or minus standard error of the mean in Figure 4 and in Supplemental Figures 7–10. These parameters are summarized in Table 1.

**Figure 4:**
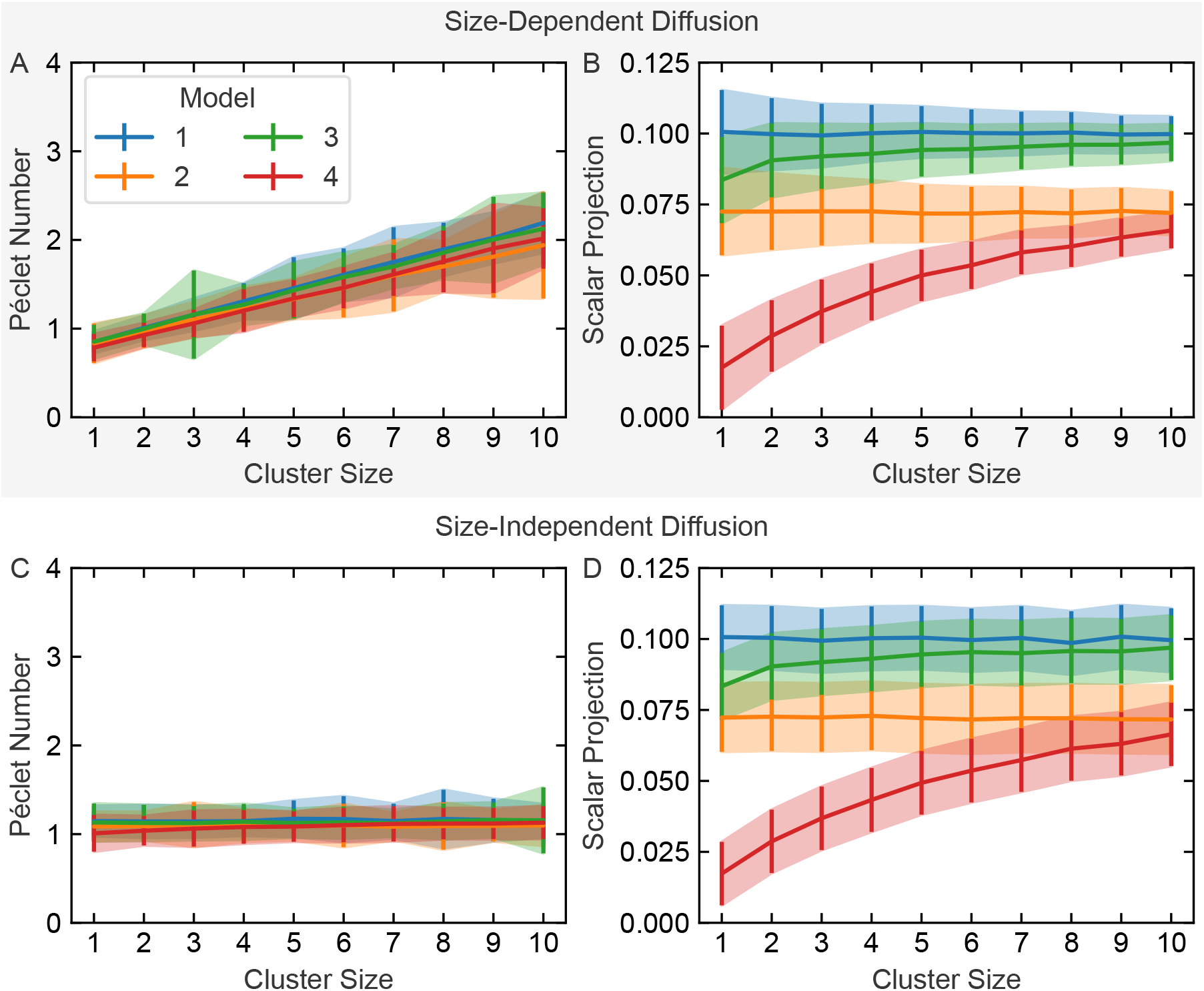
Comparison of models via Péclet number (left column) and scalar projection (right column) for *N* = 100 simulated cells with size-dependent (top row) and size-independent diffusion (bottom row). Colors distinguish models; lines with error bars and shaded area shown mean and standard error of the mean for each output (Péclet number and scalar projection) obtained through MC sampling as in Fig 3. A and C: Size-dependent diffusion is necessary to observe an approximately linear increase in Péclet number as a function of cluster size. Distinguishing models 1 through 4 based on Péclet number does not appear possible given the overlapping measurements. B and D: Scalar projection of the cluster displacement onto the cortical flow displacement can be used to distinguish the models. Model 1: the scalar projection matches the flow; Model 2: the scalar projection is reduced by the average fraction of time spent bound to the flowing cortex but does not depend on cluster size; Model 3: the scalar projection saturates to the scalar projections in Model 1 from Model 2; and Model 4: the scalar projection saturates to a much lower value than in Model 3.

#### 2.6 Code Availability

We used Python 3.8.7 (Python Software Foundation, https://www.python.org/), Matplotlib (Hunter, 2007), Pandas (McKinney, 2010), NumPy (Harris et al., 2020), SciPy (Virtanen et al., 2020), Numba (Lam et al., 2015), seaborn (Waskom, 2021), Joblib (https://joblib.readthedocs.io), and Jupyter (Pérez and Granger, 2007) for our simulations and analysis. The code used to make the figures is available as an GitHub repository at https://github.com/zmurchok/par-protein-transport-model.

### 3 Results

#### 3.1 All Models Match Experimentally Observed Trend in Péclet Number

In order to constrain the parameter space to biologically realistic values, we first preformed a parameter sweep for Models 1–4 (Section 2.5) to determine parameter ranges that are consistent with the cluster size dependent Péclet numbers observed in Dickinson et al. (2017). Using these parameter ranges, we generated *N* = 100 unique parameter sets, corresponding to 100 cells. Next, using the MC sampling method, we obtained 500 displacements for each cluster size 1–12 for each parameter set, and calculated the Péclet number for each displacement. To compare the resulting displacement distributions with protein cluster transport observed in experiments (Dickinson et al., 2017), we binned the clusters by size, and plotted the Péclet number distributions for clusters of size 1–3, 4–6, 7–9, and 10–12 using boxplots in Figure 3.

The Péclet number distributions for models 1–4 all agree with experimental data (since parameters were calibrated to this data). First, note that the Péclet number increases with cluster size across all models and across a wide range of parameters. This confirms the main hypotheses from the literature, namely, that there exists a size-dependent effect ensuring that larger protein clusters are more easily transported by cortical flow. Moreover, we estimated the mean Péclet number obtained from experimental observations from Figure 6H of Dickinson et al. (2017) and show these data in Figure 3 as black points connected with a dotted line. Our simulated clusters (boxplots) match this observed linear increase in mean Péclet data.

The linear increase in mean Péclet number critically depends on the fact that the diffusion coefficient depends on cluster size through the drag coefficient *γ* (or *γ*_*c*_ in model 4). Recall that as clusters grow in size they experience more drag from the environment and thus have a smaller overall diffusion coefficient *D*_*n*_ (equation (2.12)). To determine the role that this size-dependent diffusion contributes to the observed increase in Péclet number, we repeated the simulation as in Figure 3 but fixed the diffusion coefficient *D*_*n*_ = *D*_3_ for all cluster sizes *n*. In this case, the distributions of Péclet number and across models does not appear to increase with cluster size. To demonstrate this, we plotted the mean Péclet number versus cluster size in Figure 4 with error bars illustrating the standard error of the mean. Figure 4A replicates Figure 3 with a different plotting convention for comparison. Figure 4C, demonstrates that when size dependent diffusion is removed, the mean Péclet number does not increase when the diffusion coefficient is fixed. We thus conclude that the Péclet number effect is driven by size-dependent diffusion.

In conclusion, Models 1-4 are all consistent with the Péclet number observations from Dickinson et al. (2017) for a range of parameters. Unfortunately this type of quantification cannot alone be used to distinguish these models. The central reason for this is that there are two size dependencies in these models: (1) size-dependent diffusion and (2) size-dependent drift. The size dependence of drift varies across models. However Péclet number quantifies that drift relative to diffusion, and the size dependence of diffusion, which is the same for all models, appears to overwhelm this metric. You cannot see the signal through the noise with this metric. We address this limitation in the next section.

#### 3.2 Distinguishing Models 1–4

Given that the observed cluster transport characteristics (Péclet number) match between all of the models considered here (Figure 4A, we sought a metric that could distinguish the models. Motivated by the observation that the observed change in Péclet number is driven by size-dependent diffusion, we propose a different metric, the scalar projection of the cluster displacement on to local cortical flow displacement over the length of time Δ*t*. We illustrate how this metric is calculated in Figure 2A. Unlike the Péclet number, the scalar projection of the displacement on to the cortical flow is not dominated by the diffusive process in the cluster transport. Instead of comparing drift to diffusion, it compares drift to the cortical flow directly, which should be experimentally accessible from data since Dickinson et al. (2017) and others use particle image velocimetry to determine the cortical flow velocity at each location in the cell.

We plot the mean scalar projection (and standard error of the mean) for the four models across cluster sizes in Figure 4B. Unlike the Péclet number, the mean scalar projection separates into four distinct size-dependent curves. This suggests that it may be possible to distinguish the models by using this metric. We observe that the scalar projection for model 1 is constant with cluster size and that the standard error decreases with cluster size. This agrees with our intuition. In model 1, the clusters always move with the cortical velocity, so their scalar projection should be equal to the cortical flow speed (0.1). Larger clusters are less affected by noise compared to small clusters since they experience more drag from the environment, so we expect the estimate of the mean scalar projection to be more precise. In model 2, the scalar projection is independent of cluster size and takes on values less than the cortical flow speed. This is reasonable since in this model, the clusters stochastically bind and unbind to the cortex and only drift with the cortex when bound. In model 3 and 4, we observe a size-dependent saturation in the mean scalar projection. This dependence is qualitatively different between the two models. In model 3, the scalar projection saturates fairly quickly near the cortical velocity since a sufficient number of binding sites leads to a large fraction of time spent bound. In model 4, on the other hand, the drag cross section of the cluster grows more slowly with size and thus the scalar projection does not reach the cortical velocity for even the largest sizes considered. Finally, note that the scalar projection is less sensitive to the size-dependent diffusion coefficient (compare Figure 4B and D).

In conclusion, the scalar projection appears to have more power to distinguish these hypotheses than the Péclet number. The essential reason is that the Péclet number compares drift to diffusion and all models have a substantial and similar size-dependent diffusion. The scalar projection on the other hand compares drift to cortical flow, which circumvents this issue and focuses on how the interaction between clusters and the cortex influence drift.

#### 3.3 Estimating Scalar Projection Across Cluster Size and Parameter Space

To further evaluate how the mean scalar projection depends on cluster size, model, and parameter choices, we analytically approximate the mean scalar projection to simplify its characterization as a function of parameters. To do so, we exploited the fact that the cluster displacements are random variables following a distribution given by equation (2.23), and the fact that the scalar projection is simply a function of that random variable. Recall that the cluster displacements of size *n* and binding state *b* are random variables that are normally distributed according to

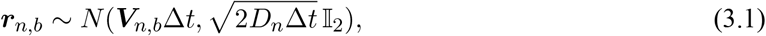

and that the scalar projection for these cluster displacements is therefore

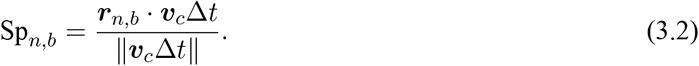

In models 1 and 4 where binding and unbinding to the cortex is not considered, it is straightforward to determine an expression for the mean scalar projection since the mean of the normal distribution is easily found as ***υ***_c_Δ*t* (model 1) or 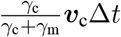 (model 4):

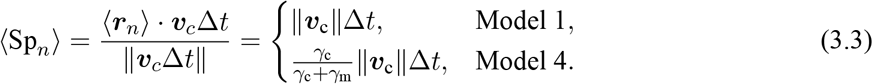

Here, ⟨−⟩ indicates the expectation of the random variable. Note that the fraction 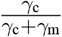 depends on the cluster size *n* through the drag coefficient *γ*_c_.

To find an expression for the mean scalar projection for clusters of size *n* in models 2 and 3, it is necessary to also average over the binding and unbinding process. Recall that the clusters in model 2 and 3 move with the cortical velocity when bound and otherwise freely diffuse, and that the binding and unbinding rates are constant in model 2 but depend on the cluster size *n* in model 3. Using a quasi-steady approximation (valid for sufficiently fast binding and unbinding rates or for sufficiently long simulation time *T*), we find that 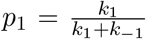 is the fraction of time spent bound, and *p*_−1_ = 1 − *p*_1_ is the fraction of time spent unbound (see Section 2.1). Thus, to find the expected scalar projection for clusters of size *n*, we first average over the bound and unbound states:

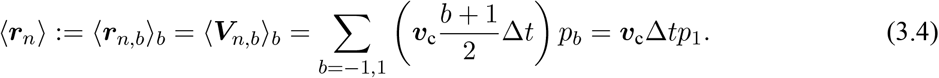

This matches our intuition: the average cluster displacement over a time Δ*t* should be equal to the displacement generated by the flowing cortex over that time interval, Δ*t*, scaled by the amount of time spent bound to the cortex, *p*_1_. Thus, for models 2 and 3, we find that

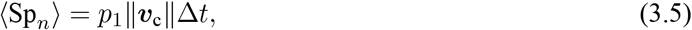

where the fraction of time spent bound, *p*_1_ is fixed for model 2 and depends on cluster size for model 3.

These theoretical estimates for the mean scalar projection agree with the mean scalar projection obtained from MC sampling. In Figure 5A, we overlaid the theoretical estimates (dashed lines) on the mean scalar projection obtained from MC sampling (an exact copy of Figure 4B). In order to make this comparison, we first calculated the theoretical estimate of the mean scalar projection for each parameter set used in the MC sampling, and then took the average over the parameter sets. We observe excellent agreement between this theoretical estimate and the simulated cells. This analytical approach will likely simplify analysis of data using this scalar projection approach since it allows for a more direct observation of how parameters, many of which are not directly estimable, would influence this metric.

**Figure 5:**
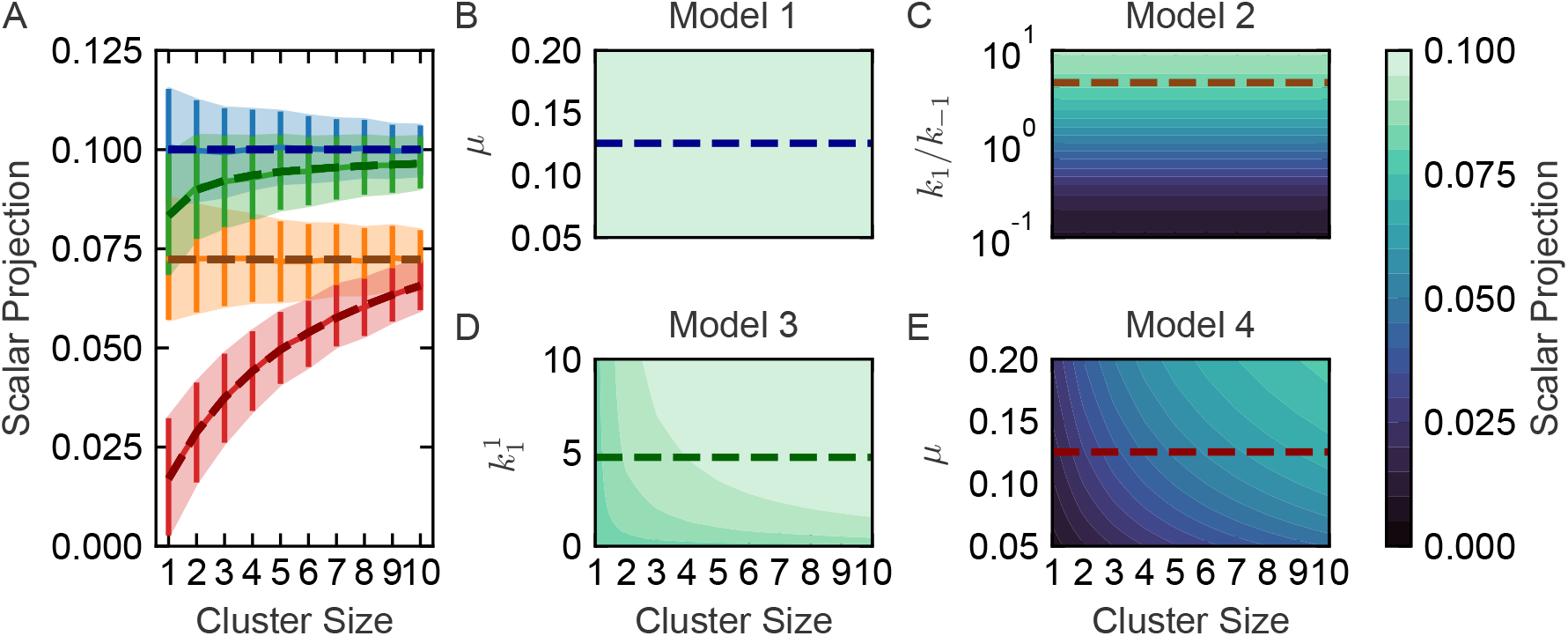
Theoretical predictions of mean scalar projection for each cluster size, the four models and parameter space. A. Dashed lines show how the predicted mean scalar projection values agree with the scalar projection values obtained through MC sampling as in Figure 4B. B-D: Mean scalar projection values as a function of cluster size and key model parameters. B. Model 1 mean scalar projection is constant and exactly matches the cortical flow speed. C. Model 2 mean scalar projection does not depend on cluster size and increases with *k*_1_/*k*_−1_. D. Model 3 mean scalar projection depends nonlinearly on cluster size and 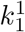 and rapidly approaches the cortical flow speed for medium sized clusters and intermediate values of 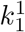 (*k*_1_/*k*_−1_ = 5 is fixed). E. Model 4 mean scalar projection depends nonlinearly on cluster size and viscosity *μ*; but remains much smaller than the cortical flow speed unless the cluster size and drag are both large.

The advantage of the theoretical estimates of the mean scalar projection is that we now can understand how the scalar projection depends on the model parameters without relying on simulations. In Figure 5B–E, we show heat maps of the mean scalar projection estimated by theory across cluster sizes and parameter spaces. Each panel illustrates the mean scalar projection for a different model, and the dashed lines correspond to the average parameter value used in the *N* = 100 parameter sets in the MC sampling method. We find that the mean scalar projection is constant and matches the cortical flow speed regardless of model parameters in model 1 (panel B); increases with the ratio *k*_1_/*k*_−1_ but is constant with respect to cluster size in model 2 (panel C); quickly saturates with respect to cluster size to the cortical flow speed in model 3 (panel D); and slowly saturates with respect to cluster size to approximately 75% of the cortical flow speed in model 4.

### 4 Discussion

In this study, we developed a model to study how PAR-3 protein clusters interact with the flowing actin cortex in single cell *C. elegans* embryos. This discrete, agent-based model (ABM) incorporates cluster size dynamics, cortex-binding and unbinding dynamics, as well as a physics-based model for the cluster’s interaction with the cortical flow. The purpose of this model is to use it as a platform to test different hypotheses for how these protein clusters interact with the cortex, and how those interactions influence their transport. Toward this end, we encoded four hypotheses for how the clusters may interact with the flowing actin cortex and tested each model against experimental observations. The models encoded were (1) that clusters are directly coupled to the cortex and moved with its velocities; (2) that clusters can stochastically bind and unbind from the cortex, independent of their size; (3) that clusters can bind and unbind from the cortex in a size-dependent fashion; and (4) that the cluster experiences a drag force from the flowing cluster.

The primary data that both motivated this study and which we use to test these hypotheses is particle tracking data from Dickinson et al. (2017). There, the authors quantify intensity of PAR3 clusters (a proxy for cluster size), track their positions over time, and calculate the Péclet number of clusters of different intensities (i.e., sizes). Based on this data, we began this study with the broad hypothesis that clusters of different sizes interact with the cortex in different ways, and that by comparing these models to this size dependent data we could elucidate the form of that cluster-cortex interaction.

As a first step, we demonstrate that each model variant (encoding the four hypotheses) could be calibrated with biophysically reasonable parameters to match the size-dependent Péclet observations. On one hand, this calibration allowed us to restrict the parameters of each model to biophysically relevant ranges. On the other, however, it demonstrates that the Péclet number is an insufficient measure to distinguish these different models. The essential reason for this is simple in retrospect. The Péclet number quantifies the relative importance of advection compared to diffusion in a system. In these models, cluster size alters both advection (which we are interested in) and diffusion (which we are not). In the regime of behavior of this system, size-dependent diffusion effects (i.e., larger clusters diffuse less) dominate the Péclet number’s size dependence, making it unsuitable for our purpose.

In response to this observation, we sought to find a way to differentiate these hypotheses based on the type of particle tracking data that is available. Our results show that a “scalar projection” measure can distinguish between the four different hypotheses. This scalar projection quantifies the level of advection of clusters of different size relative to the cortical flow. Each of these models produces a distinct size-dependent scalar projection curve that match biological intuition. For this reason, we suggest that particle tracking data could be analyzed in conjunction with particle image velocimetry (to measure the cortical flow) to facilitate the calculation of scalar projection.

This is of course not the only approach to differentiating these models. Techniques to either directly or indirectly observe the cluster’s state (size and binding status) from single particle data (Du and Kou, 2020; Falcao and Coombs, 2020; Kowalek et al., 2019; Mellnik et al., 2016; Menssen and Mani, 2019; Robin et al., 2014) would be a more direct way to test this. Experimental perturbations to cortical flow (Mittasch et al., 2018) may also be useful in distinguishing models. Alternatively, engineered clusters could be used to study transport properties. Recent experimental work by Chang and Dickinson (2021) took this approach. Their new data suggests that clusters of size 3 or larger indeed exhibit directed transport, that the transport of larger clusters is more ballistic (as opposed to diffusive), and that the cluster diffusion constant decreases with cluster size. While this demonstrates clear size dependent effects on transport, we note that both Models 3 and 4 exhibit these basic characteristics (larger clusters move more ballistically and with lower diffusion constant). Thus it remains unclear whether these size-dependent transport effects result from mainly drag effects or more direct binding to the flowing cortex. Further analysis of this or similar data with the aforementioned scalar projection metric may help resolve this.

We also developed new approximate analysis methods to facilitate the above investigation. Full stochastic simulation of these models is relatively straightforward; however, such simulations can be computationally slow and creates sampling issues since there are far fewer instances of large clusters in the model than smaller ones. In particular, we found that long simulations were required to produce enough data to construct reliable quantification for rarer, larger clusters, dramatically oversampling data for smaller clusters in the process (note that Dickinson et al. (2017) found that most PAR-3 clusters were small in experimental data). Since both Péclet number and scalar projection quantification only use displacement data rather than positional data, we analytically constructed approximate probability distributions for displacements of clusters of each size for each model. This greatly simplifies simulation of Péclet number and scalar projections since we can sample displacements from the distribution for each cluster size independently and construct these measures directly. This approach also allowed us to formulate semi-analytical predictions for the size dependent mean scalar projection for each model, eliminating the need for simulations. This speeds analysis of the effects of parameters on predictions for each model and should simplify any further analysis of these models based on new data or analysis in the future.

While there are several other studies that seek to understand how the dynamics of protein clustering in *C. elegans* and other systems affects their movement and biology (Agudo-Canalejo et al., 2020; Bergeler and Frey, 2018; Dawes and Munro, 2011; Gambin et al., 2006; Knight et al., 2010; Kober et al., 2019; Liu et al., 2020; Sailer et al., 2015; Yi et al., 2012; Ziemba and Falke, 2013) our study is the first to explicitly and model the transport of protein clusters during early embryogenesis in *C. elegans*. Nonetheless, there are several limitations of our study. First, we did not model membrane binding and unbinding in this study and chose to focus on cortex-cluster binding. This is due to the lack of information about the biophysics at play. For example, it is not clear weather clusters that unbind from the membrane remain visible in the imaging assay or what happens to those clusters once disassociated. This could however readily be added to the model in the future. A second limitation is that we have ignored the position dynamics and cell geometry by focusing only on the cluster displacements instead of the cluster’s movement through the cell (Dickinson et al. (2017) quantified track lengths for clusters of various). A third limitation is that there are many feedback loops between PAR-3 clustering and other proteins which we have ignored. During the polarization of the embryo, PAR-3 proteins promote local association of PAR-6/PKC-3 with active CDC-42 GTPase (Kravtsova and Dawes, 2014; Lang and Munro, 2017; Sailer et al., 2015; Seirin-Lee et al., 2020). This process may alter the transport of PAR-3 clusters as it binds and unbinds with other proteins or modulates the local flow of the cortex through GTPase signaling (which affects the cytoskeleton). More detailed models, constrained by experimental data, could be used to address these limitations, but would stray from the main intent to study cluster-cortex interactions.

Overall, we developed a discrete, agent-based model to study how PAR-3 protein clusters interact with the flowing actin cortex in the context or early embryogenesis and used this model to test four biophysical hypotheses for how the clusters may interact with the cortex. While we found that all models can agree with existing experimental observations, we proposed an alternate metric for analysis that can distinguish the hypotheses from sufficiently detailed single particle tracking data. Our study presents the first computational steps in unraveling how biophysical interactions between protein clusters and cortical actin flow coordinate cluster transport.

## 5 Acknowledgments

We acknowledge John J. Vastola for helpful discussions. This work was supported by a National Science Foundation, United States, grant DMS1562078 (to WRH) and a Natural Sciences and Engineering Research Council of Canada (NSERC) Postdoctoral Fellowship Award (to CZ).

