## Supplemental Information for "Biophysical Models of PAR Cluster Transport by Cortical Flow in *C. elegans* Early Embryogenesis"


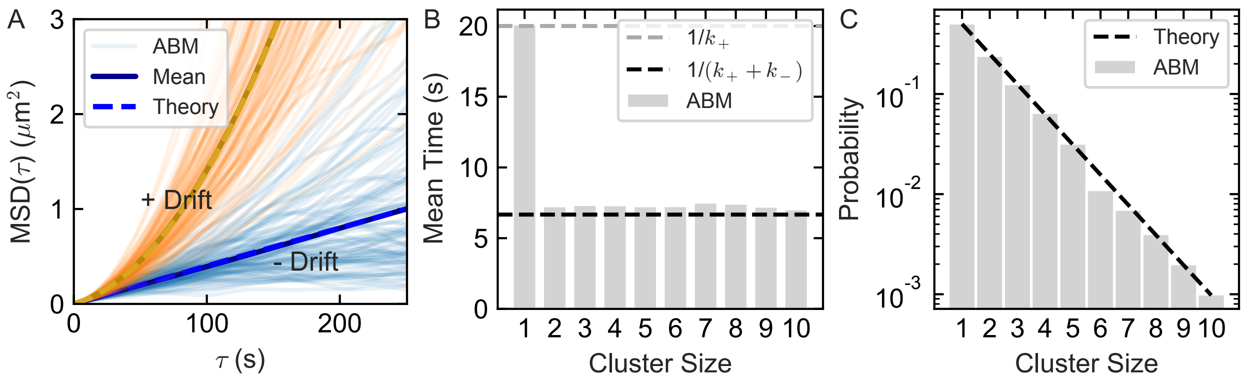


**Supplemental Figure 1:** Validation of ABM simulation with size-independent diffusion. We simulated 1000 clusters without drift ($v = 0$, "- Drift") and 1000 clusters with drift ($v = 0.01$, "+ Drift") with $D = 0.001$. A. Averaged mean-square displacement (MSD($\tau$) $\mu$m$^2$) over all simulated clusters as a function of time-lag $\tau$ s (solid lines) agrees with the MSD expected from theory (dashed lines). B. Mean time in each cluster size state obtained from ABM simulations agrees with that predicted by theory. C. Probability that a cluster is in size $n$ matches that predicted by theory for $k_+ = 0.05$ and $k_- = 0.1$.



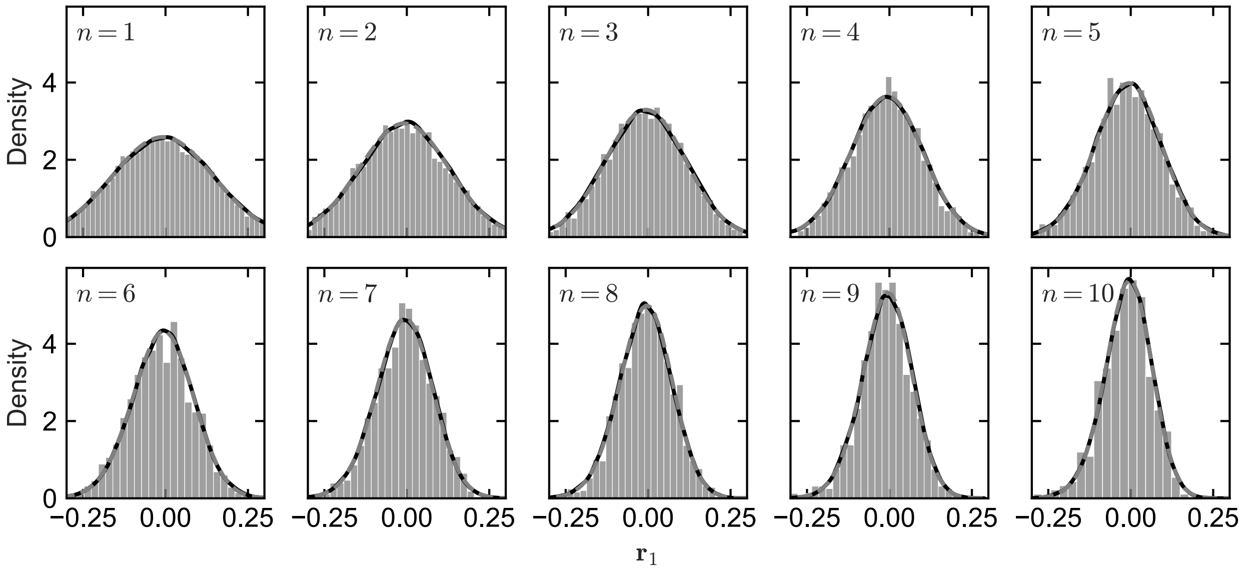


**Supplemental Figure 2:** Probability density distributions of the $x$-component of cluster displacements, $\boldsymbol{r}_1$, obtained from the simulations of the agent-based model (ABM; histograms) match the kernel density estimates of distributions obtained from Monte-Carlo (MC) sampling (solid lines) from quasi-steady distributions (dashed lines) across clusters of size $n = 1, \ldots 10$ in Model 2 with parameters as in Figure 2.

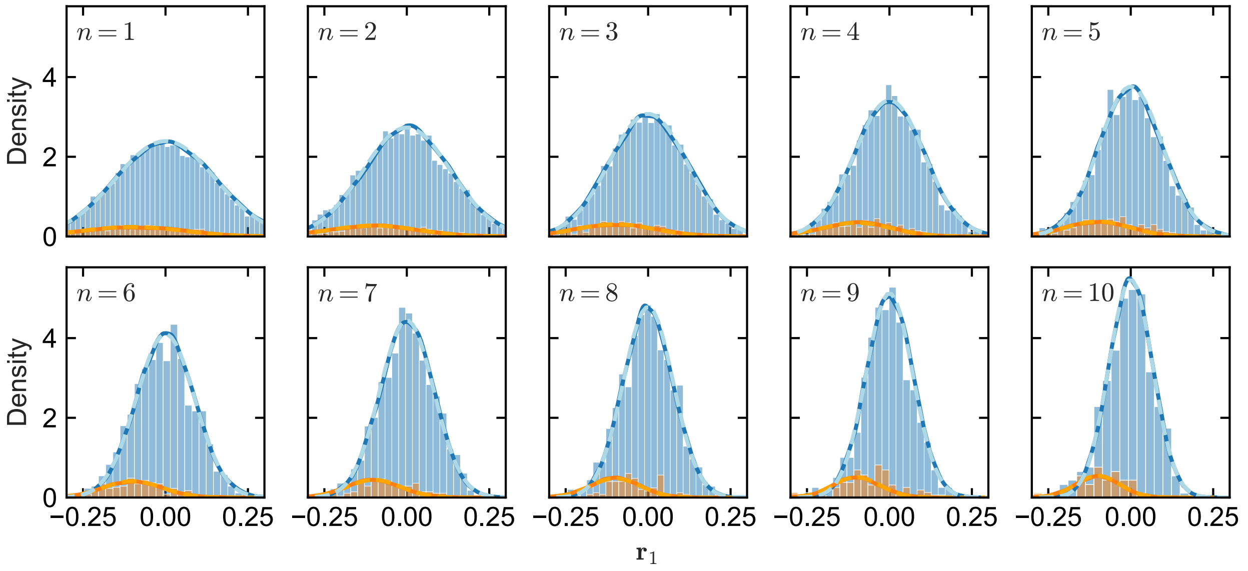


**Supplemental Figure 3:** Probability density distributions of the $x$-component of cluster displacements, $\boldsymbol{r}_1$, obtained for Model 2, as in Supplemental Figure 2, except the distributions are split into bound (orange) and unbound (blue) states. Distributions obtained from ABM simulations (histograms), MC sampling (solid lines), and theory (dashed lines) match with each other.

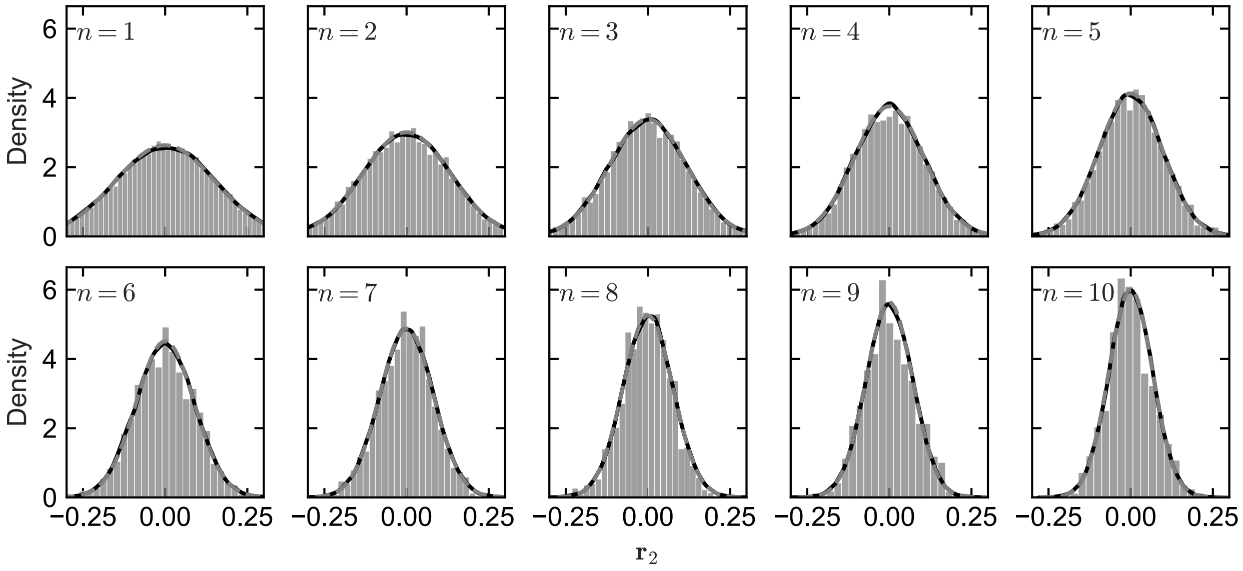


**Supplemental Figure 4:** Probability density distributions of the $y$-component of displacements obtained for Model 2, as in Supplemental Figure 2. Distributions obtained from ABM simulations (histograms), MC sampling (solid lines), and theory (dashed lines) match with each other.

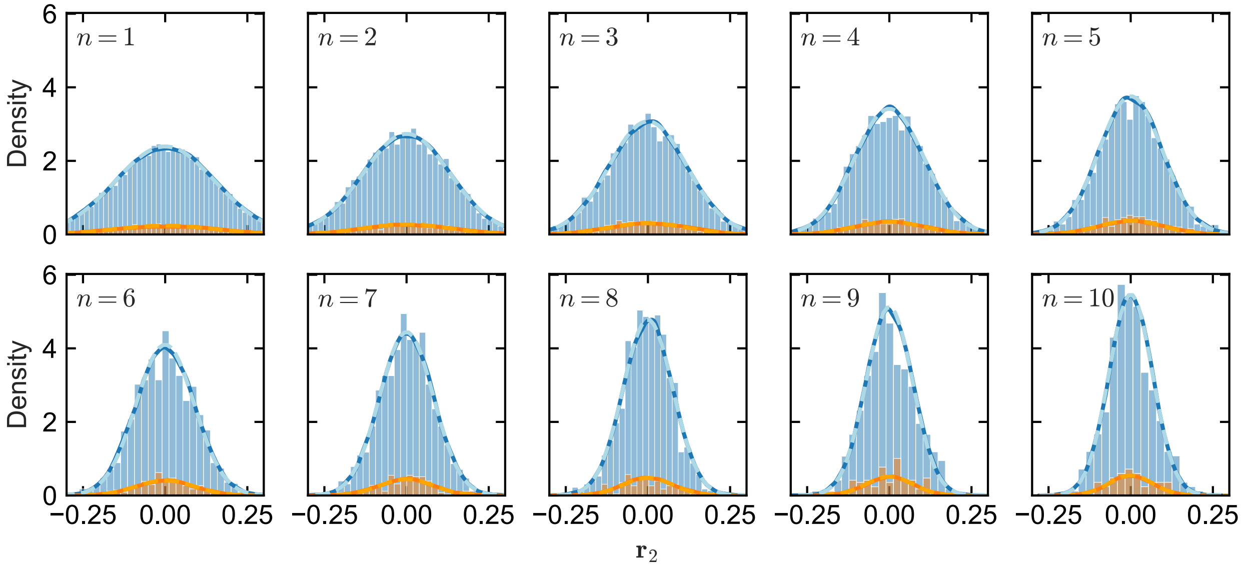


**Supplemental Figure 5:** Probability density distributions of the $y$-component of displacements obtained for Model 2, as in Supplemental Figure 5, except the distributions are split into (bound) orange and unbound (blue) states. Distributions obtained from ABM simulations (histograms), MC sampling (solid lines), and theory (dashed lines) match with each other.

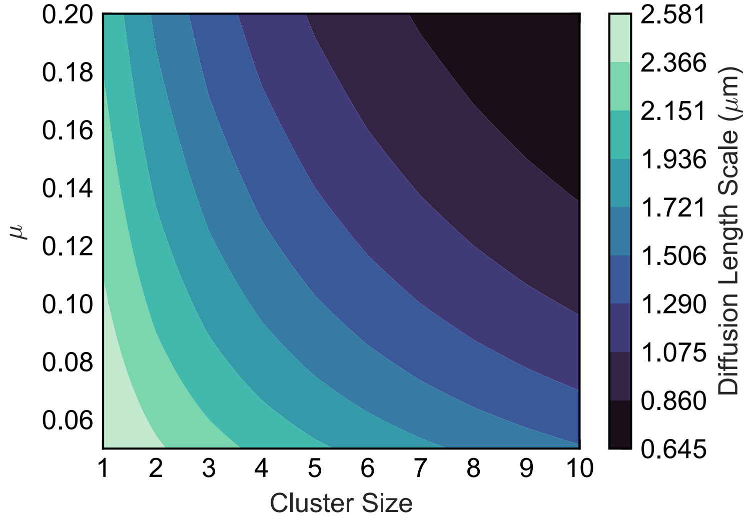


**Supplemental Figure 6:** Estimated diffusion length scale $L = \sqrt{D_n T}$ $\mu$m for freely diffusing clusters of size $n$ and total time $T = 500$ s with $\sigma = 0.01$.

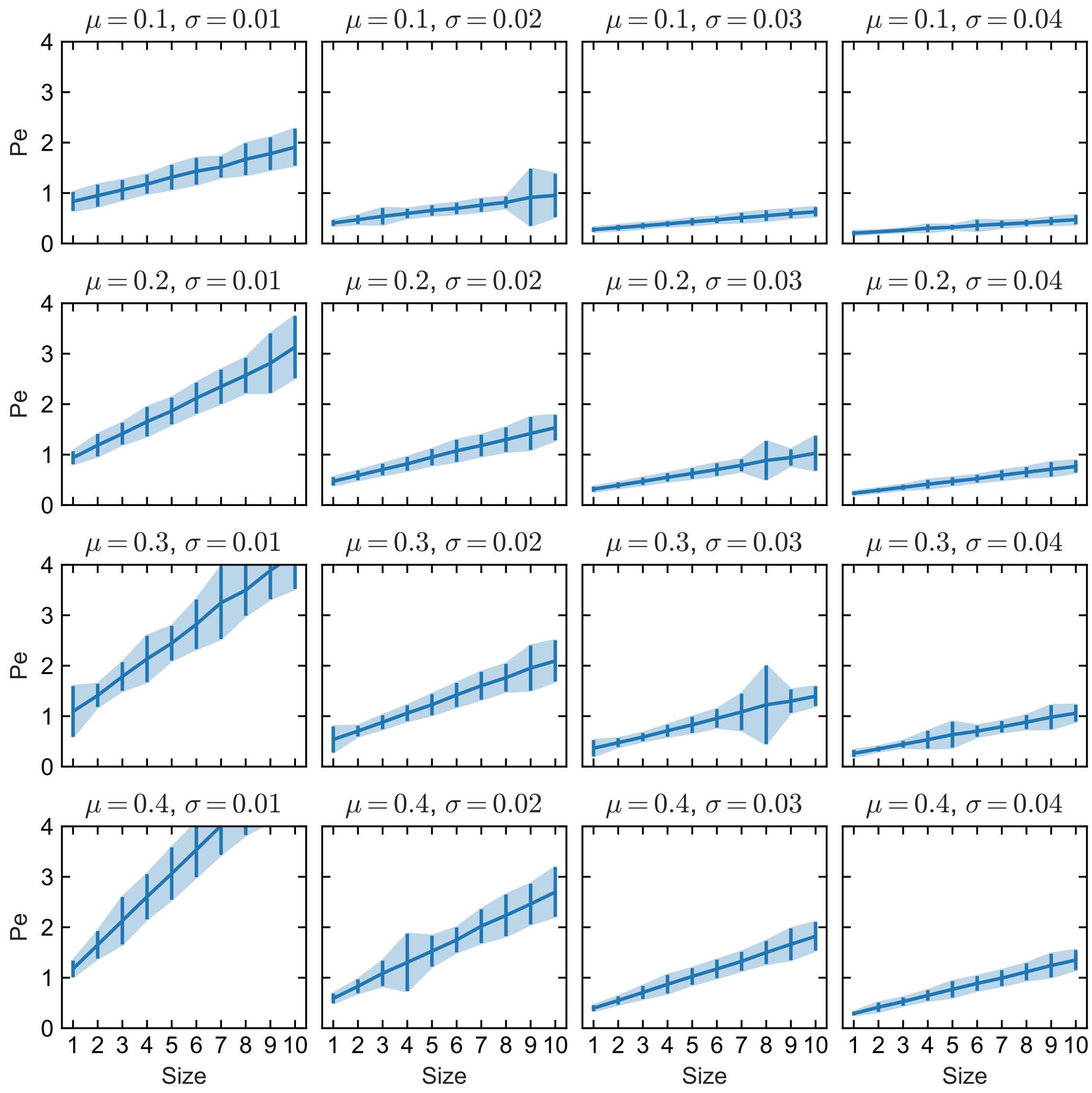


**Supplemental Figure 7:** Model 1 parameter sweep. We sampled 500 displacements for each cluster size $n = 1, \ldots, 10$ from $N = 100$ cells with the parameters indicated as the title of each panel, and show the mean Péclet number as a function of cluster size for each parameter set (error bars show standard error of the mean). Increasing the viscosity (moving down panels) reduces the effect of diffusion, thus increasing the Péclet number. Increasing the noise level (moving right panels) increases the effect of diffusion, thus decreasing the Péclet number. Note that all panels share the same $y$-axis thus some data is not shown (e.g., for large $\mu$ and small $\sigma$).

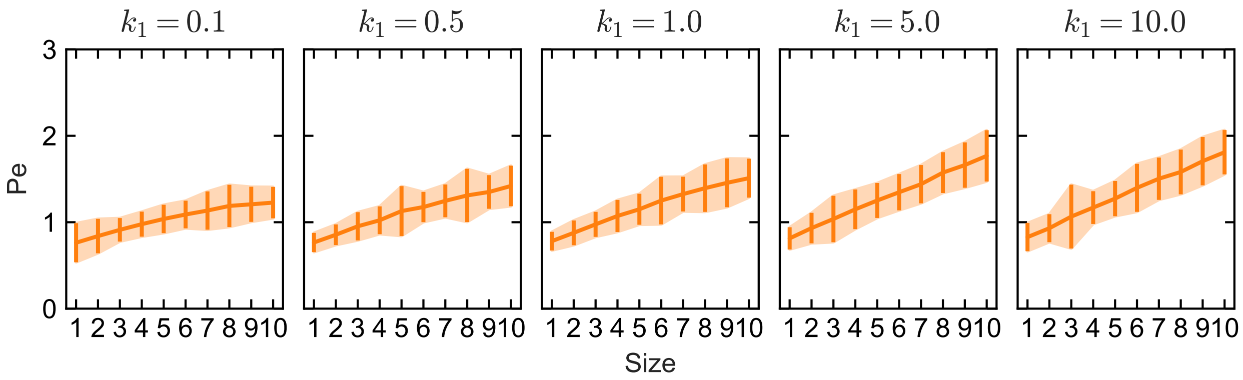


**Supplemental Figure 8:** Model 2 parameter sweep. We sampled 500 displacements for each cluster size $n = 1, \ldots, 10$ from $N = 100$ cells with the parameters indicated as the title of each panel with $k_{-1} = 1$, $\mu = 0.1$, and $\sigma = 0.01$. We show the mean Péclet number as a function of cluster size for each parameter set (error bars show standard error of the mean). Increasing $k_1$ (relative to $k_{-1}$) increases the time spent bound to the flowing cortex, thus increasing the Péclet number.

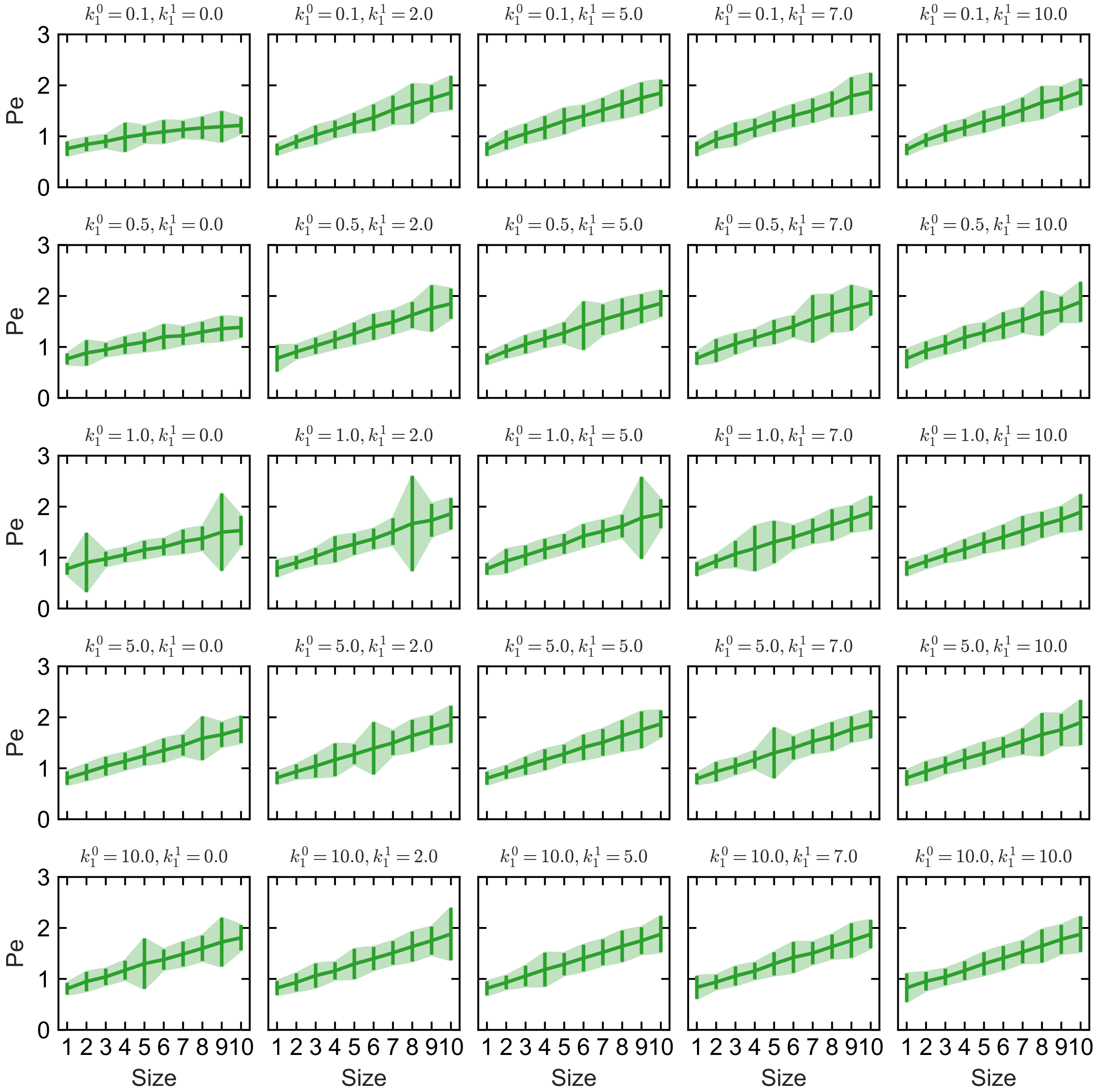


**Supplemental Figure 9:** Model 3 parameter sweep. We sampled 500 displacements for each cluster size $n = 1, \ldots, 10$ from $N = 100$ cells with the parameters indicated as the title of each panel with $k_{-1} = 1$, $\mu = 0.1$, and $\sigma = 0.01$. We show the mean Péclet number as a function of cluster size for each parameter set (error bars show standard error of the mean). Increasing $k_1^0$ (relative to $k_{-1}$) increases the time spent bound to the flowing cortex, thus increasing the Péclet number. Increasing $k_1^1$ also increases the Péclet number provided that $k_1^0$ is small (relative to $k_{-1}$).

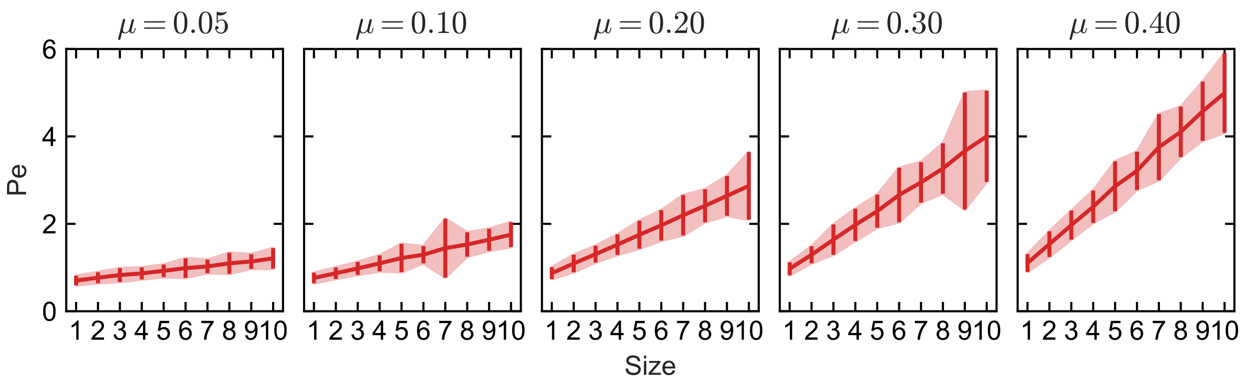


**Supplemental Figure 10:** Model 4 parameter sweep. We sampled 500 displacements for each cluster size $n = 1, \ldots, 10$ from $N = 100$ cells with the parameters indicated as the title of each panel with $\sigma = 0.01$. We show the mean Péclet number as a function of cluster size for each parameter set (error bars show standard error of the mean). Increasing $\mu$ increases the the Péclet number.

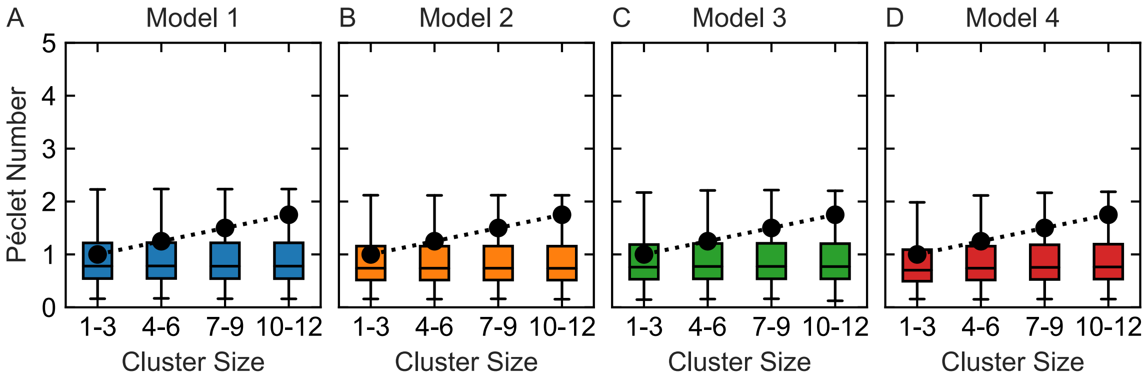


**Supplemental Figure 11:** Models with size-independent diffusion. The diffusion coefficient does not depend on cluster size (here $D_n = D_3$ for all $n$). Black points show mean Péclet numbers estimated from experimental observations of cluster transport from Figure 6H of Dickinson et al. (2017), while boxplots illustrate simulated Péclet number distributions obtained from MC sampling when cluster displacements are grouped into four size categories. Parameters are exactly as in Figure 3.
